# Bacterial major vault protein homologs form divergent vault structures

**DOI:** 10.64898/2026.11.06.764734

**Authors:** Jake C. Lachowicz, Mohammed Hoque, Anbo Zhou, Tong Zhang, Matthew C. Franklin, Richard Copin, Katherine D. Cygnar, Andrew J. Murphy, Kei Saotome, Andrew D. Baik

## Abstract

Vaults are large, enigmatic macromolecular protein complexes composed of symmetrically arranged major vault protein (MVP) subunits with a wide variety of proposed functions. While MVP is conserved in most eukaryotes, its prevalence in other domains of life has not been well studied. Furthermore, whether bacterial MVP homologs assemble into vaults remains unknown. Here, we identify and examine a subset of bacterial MVP homologs and report cryoEM structures of an intact vault from *Corallococcus praedator* and a half-vault from a Deltaproteobacteria bacterium. Both bacterial vault complexes exhibit architectures unlike any other previously characterized vault with barrel/cap symmetries of D35/C13 and C48/C16, respectively. These data demonstrate that bacterial MVP homologs can assemble into vault structures and highlight the structural diversity of vaults across domains of life.

## Introduction

The vault is a large 13 MDa macromolecular protein complex first discovered serendipitously by Kedersha and Rome in 1986^1,2^. However, after more than 40 years since its discovery, a complete understanding of the vault’s evolutionary history, structural diversity, and functional roles remains elusive. High resolution structures of vaults are limited to mammalian orthologs from *Homo sapiens* (Human)^3–6^, *Rattus norvegicus* (Rat)^7–11^, and *Oryctolagus cuniculus* (Rabbit)^12^, all of which have virtually identical architectures. Mammalian vaults are composed of 78 protomers of major vault protein (MVP), each containing 9 anti-parallel β-sheet repeat domains, an SPFH (Stomatin, Prohibitin, Flotillin, and HflK/C) shoulder domain, a cap-helix domain, and a cap-ring domain^6,12,13^. The D39-symemtric mammalian vault complex is composed of two C39-symmetric half-vaults, each breaking into C13 symmetry at their cap-ring domains^6,12^. Although assembled as an intact vault^14^, vaults can open to form two half-vaults^15^, and have even been shown to undergo half-vault exchange^16^.

Vault/MVP has been described to have roles in diverse cellular processes such as autophagy^17^, signal transduction^18–21^, immunity^22–25^, cellular transport^26,27^, and even cancer progression^28^. However, how these reported functions are interconnected and how non-MVP proteins associated with vaults (minor vault proteins) contribute has not been fully established. Notably, the large, hollow morphology of vaults and presumed low immunogenicity have also attracted studies into their use as delivery vehicles for therapeutic cargo^29–34^.

MVP homologs are broadly distributed within most eukaryotic super-groups but are noticeably absent in plants, fungi, and some model organisms including *Saccharomyces cerevisiae* and *Drosophila melanogaster*^35,36^. While mammalian vaults are composed of one unique MVP monomer with 9 repeat domains, there appears to be variability in the subunit composition and domain arrangement of other eukaryotic MVPs. For example, vaults of *Dictyostelium discoideum* and within the supergroup Discoba are formed by multiple distinct MVP paralogs^37–40^, though *Dictyostelium discoideum* vaults share similar dimensions with mammalian vaults^38^. Additionally, MVP of the ameboflagellate *Naegleria gruberi* is predicted to have <9 repeat domains^41^, suggesting they could form smaller vaults than those found in mammals.

While molecular evolutionary studies suggest that MVP was present in the last eukaryotic common ancestor^35^, sequences predicted to be MVP homologs have also been found in bacterial genomes^35,42,43^. It is currently postulated that bacteria obtained MVP by horizontal gene transfer (HGT) from eukaryotes^35,42^. However, the functional role of bacterial MVP homologs and the architecture of their oligomeric assemblies remain unknown.

To gain insight into the evolution of MVP within bacteria, we performed a comparative genomic analysis of the MVP sequence space and identified novel MVP homologs in bacteria with low sequence identity to human MVP. We purified a subset of these homologs and found that several have a propensity to assemble into oligomeric architectures resembling mammalian vaults. Using single particle cryogenic electron microscopy (cryoEM), we obtained high resolution reconstructions of the intact vault from *Corallococcus praedator* (CpVault) and a half-vault from a Deltaproteobacteria bacterium (DbVault). CpVault and DbVault form superstructures with divergent size, symmetry, and cap architectures compared to other known vaults. We dissect the divergent structural properties of these bacterial vaults to gain insights into the evolution of MVP/vault structure.

## Results

### Phylogenetic analysis of the major vault protein homologs across domains of life

Previous studies identified MVP homologs only through BLASTp searches of human, rat, or various leishmania, trypanosome, or bacterial sequences^35,41,43^. To expand upon these analyses and determine the prevalence of MVP homologs throughout the domains of life, we performed a sequence- and structure-based comparative analysis. First, we identified a highly conserved stretch of nucleotides in the human MVP (hMVP) gene, which corresponded to residues 633-668 in the cap-helix of hMVP (**Extended Data Fig. 1a).** We then performed BLASTp searches using this peptide and the complete hMVP protein to form a sequence similarity-based dataset of putative MVP homologs (**Extended Data Fig. 1b**). We further performed FoldSeek searches using hMVP against the AlphaFold v6 structure database and identified *Corallococcus carmarthensis* MVP as the top bacterial hit (Uniprot: A0A3A8K409)^44–47^. After another FoldSeek search using the *Corallococcus carmarthensis* MVP sequence, we pooled the results of this search with the FoldSeek hits from hMVP to form our structural similarity-based dataset. Merging the sequence- and structure-based datasets yielded 3,959 unique putative MVP homologs, of which 1,083 (27.4%) were recovered by both approaches, 2,428 (61.3%) by BLASTp only, and 448 (11.3%) by Foldseek only (**Extended Data Fig. 1c, d**).

To map the sequence space of MVP throughout the tree of life, we constructed a cladogram from our combined MVP homolog dataset. As expected, MVP homologs are broadly distributed throughout eukaryotic lineages^35^. However, 383 (∼10%) and 22 (<1%) sequences were identified in prokaryotes and archaea, respectively (**Fig. 1a**). Similar to previous reports ^35,42,43,48^, most prokaryotic MVP homologs (205, ∼53%) belong to Myxococcia and Cyanophyceae. In archaea, the limited number of MVP homologs suggested either low prevalence or low sequencing coverage. To map the evolutionary path of bacterial MVPs, we inferred a phylogeny using sequences beyond the last repeat domain (denoted cap domain) (**Fig. 1b**). In this tree, 288 bacterial and 21 archaeal cap domains cluster into several well-defined clades. The largest of which encompasses roughly one-third of all prokaryotic sequences. This lineage branches basally with early-diverging protist groups (Discoba, SAR), rather than with Metazoa. This topology is consistent with an early evolutionary event rather than a recent acquisition, although broader sampling will be needed to distinguish among evolutionary scenarios.

**Fig. 1:**
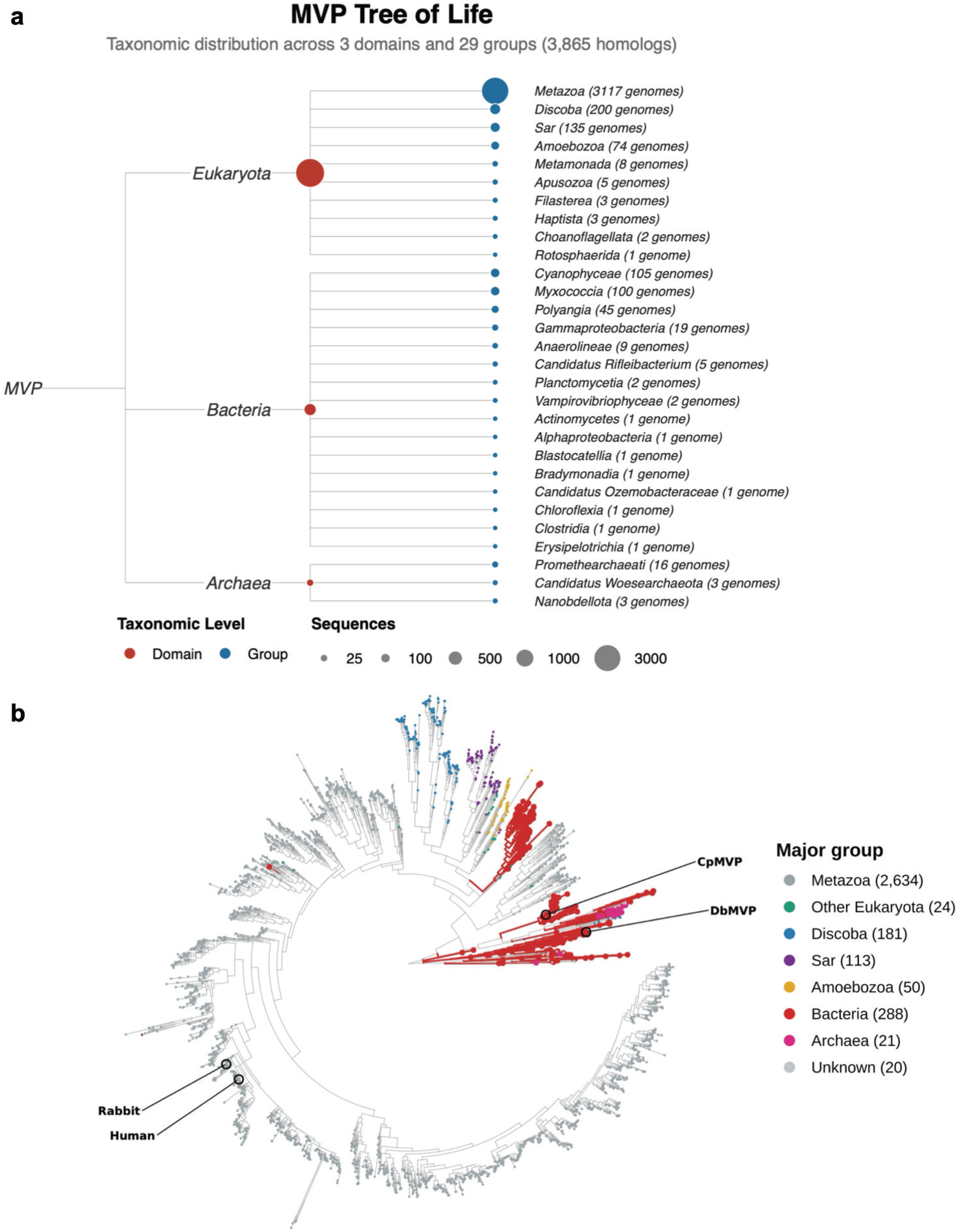
Phylogenetic analysis reveals a subset of major vault homologs in prokaryotic organisms. **a** Cladogram of MVP homologs from the three domains of life derived from 3,825 sequences from a comprehensive BlastP and Foldseek search (see materials and methods for details). **b** Maximum-likelihood phylogeny of 3,331 length-filtered MVP cap domains, inferred in IQ-TREE under the site-heterogeneous LG+C20+F+G PMSF mixture model (1,000 bootstrap and 1,000 SH-aLRT replicates), using a FastTree (LG+Γ) topology as the guide tree. The final tree is rooted with MinVar-Rooting^67^. Tips are colored by major taxonomic group (legend; counts in parentheses): Metazoa (grey), other eukaryotes, Discoba, Sar and Amoebozoa (protist lineages), and the prokaryotes Bacteria (red) and Archaea (magenta).

We further analysed our MVP homologs by sequence length (**Extended Data Fig. 2a, b**). The distribution of sequence length ranged from 31–3,204 amino acids (a.a.). 106 of these sequences are likely misannotated as they carried a foreign sequence fused to the MVP module at one or both termini. Correcting these genes (see Methods) lowered the maximum length from 3,204 to 1,400 a.a. and ∼46.2% of sequences were in the ∼850–900 a.a. range. This is in line with human (893 a.a.), rat (861 a.a.), and rabbit (890 a.a.) MVPs. Eukaryotic homologs were longer (median 856) than bacterial homologs (median 758 a.a.) on average. Interestingly, the bacterial shortening was clade-dependent: Cyanophyceae remained near eukaryotic length (median ∼872 a.a.), whereas Myxococcia were shorter (median ∼758 a.a.). Among eukaryotes, protist lineages (Amoebozoa, median ∼829 a.a.; Sar, median ∼847 a.a.) trended slightly shorter than vertebrates (865 a.a.).

### Bacterial MVP homologs from Myxococcia and Cyanophyceae form vault or vault-like structures

We selected a subset of six bacterial MVP homologs from our phylogenetic analysis for experimental characterization. Four are from the class Myxococcia: *Corallococcus praedator* MVP (CpMVP; Uniprot: A0ABX9QQB9, 32.3% identity to human MVP [see supporting information]), *Melittangium boletus* MVP (MbMVP; Uniprot: A0A250IM67, 32.4% identity), an MVP sequence annotated from Deltaproteobacterium (DbMVP; Uniprot: A0A661NLQ3, 22.4% identity), and *Corallococcus aberystwythensis* MVP (CaMVP; Uniprot: A0A3A8QLP0, 32.0% identity). Two are from the class Cyanophyceae: *Planktothrix pseudagardhii* MVP (PpMVP; Uniprot: A0A9W4G7X1, 52.77% identity) and *Okeania* sp. MVP (OsMVP; Uniprot: A0A6P0RA99, 48.4% identity). AlphaFold predicted these homologs to fold similarly to eukaryotic MVP, with each harbouring repeat domains, a characteristic SPFH shoulder domain, and a C-terminal cap-helix **(Extended Data Fig. 2c-f**; materials and methods)^44–46^. Interestingly, the number of repeat domains predicted for DbMVP (6), CpMVP (7), CaMVP (7), and MbMVP (7) are less than the 9 repeats observed in most other MVPs including human, suggesting they may assemble into vaults with unique morphology.

Each of the six MVP homologs were expressed in *E. coli*, affinity purified via C-terminal tag, and subjected to cryoEM imaging to assess vault formation and morphology (**Extended Data Fig. 3; Supplementary Fig. 1**). Micrographs of PpMVP, MbMVP, and CaMVP displayed fractured or incompletely assembled vault-like structures, while OsMVP formed eye-shaped higher-order structures that were not reminiscent of vaults (**Extended Data Fig. 4a-d**). Although PpMVP and OsMVP particles were too heterogeneous to produce interpretable 3D reconstructions, we obtained low resolution reconstructions of CaMVP (CaVault) and MbMVP (MbVault) (**Extended Data Fig. 4e, f**). These maps were similar to each other and revealed shells with morphology consistent with known vaults but with apparent breakages along the central axis, suggesting perturbed assembly. DbMVP formed homogenous dome-shaped structures that appeared to be half-vaults (**Extended Data Fig. 5a**). Micrographs of CpMVP clearly exhibited homogenous intact vault particles with the canonical barrel and cap shape of mammalian vaults (**Extended Data Fig. 5b**). The homogeneity and apparent structural integrity of DbMVP and CpMVP particles allowed us to pursue high resolution cryoEM structure determination.

### Divergent structures of the Deltaproteobacteria half-vault and *C. praedator* vault

2D classification of the DbMVP particle (DbVault) revealed only half-vaults with no intact vaults observed (**Extended Data Fig. 5a**). Top-down 2D classes and ab initio reconstruction without imposed symmetry suggested this particle was comprised of 48 protomers. 3D reconstruction with C48 symmetry imposed produced a 2.8 Å resolution half-vault map that is approximately 210 Å long and 480 Å wide, significantly shorter and wider than the 335 Å x 400 Å rabbit half-vault (**Fig. 2a**). This map contained well-resolved density for much of the protein excluding disordered loops (**Fig. 2b; Extended Data Fig. 5c; Extended Data Figure 6a**). We truncated sidechains of repeat domains R1 and R2 (residues 26-123) due to weak density, likely the result of the half-vault state. Each DbMVP protomer exhibits a canonical MVP structure that consists of 6 repeats of antiparallel β-sheets, an SPFH shoulder domain, and a cap-helix domain. The shorter cap-helix of DbMVP (∼120 a.a.) compared to rabbit MVP (∼150. a.a.), three fewer repeat domains (6 vs 9), and increased number of protomers (48 vs 39) together underlie the unique flattened shape of the DbVault half (**Fig. 2e, f, h**).

**Fig. 2:**
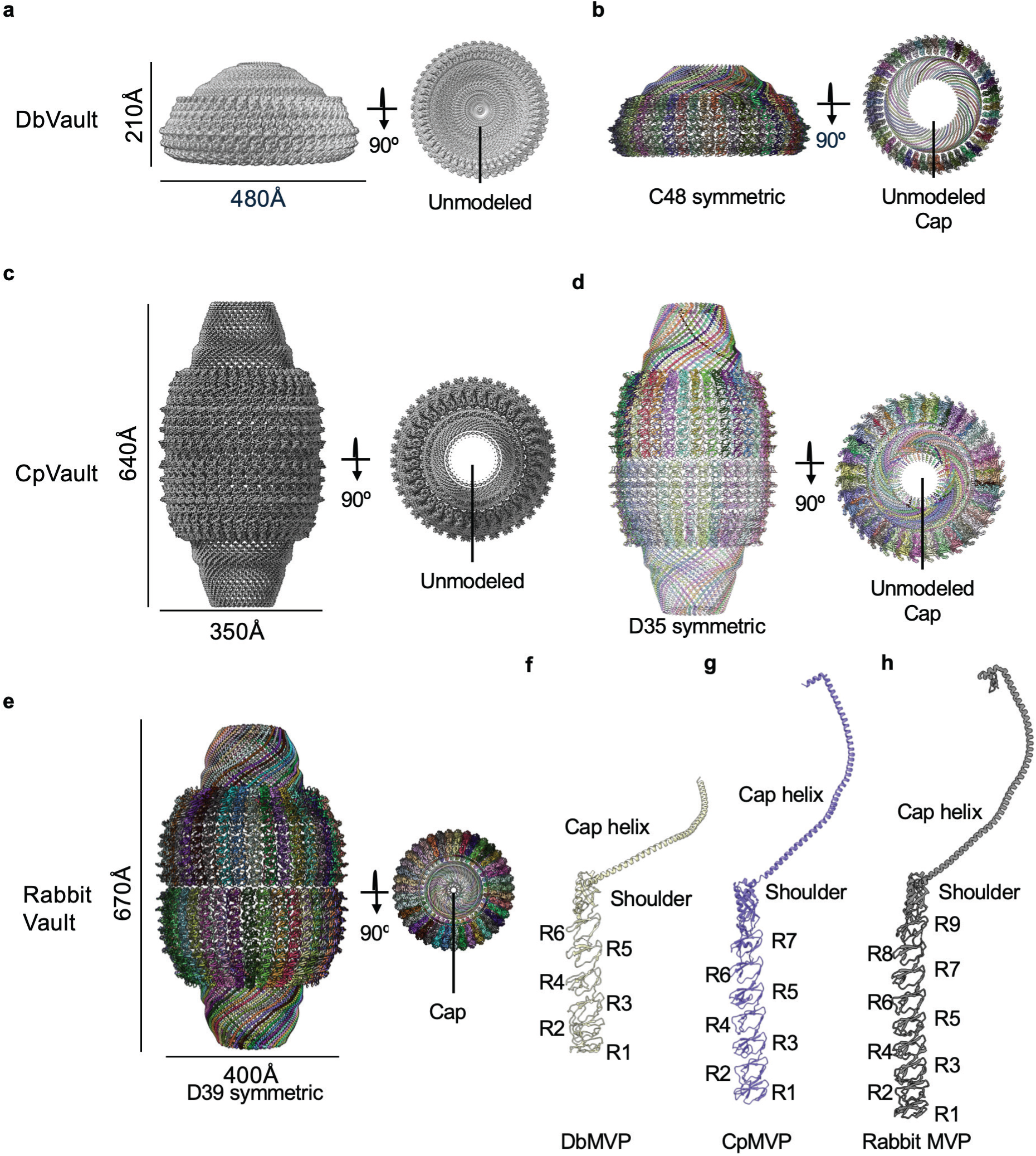
Macromolecular organization of DbVault and CpVault. **a,b** CryoEM reconstruction (**a**) and molecular model (**b**) of DbVault. **c,d** CryoEM reconstruction (**c**) and molecular model (**d**) of CpVault. **e** Molecular model of a representative mammalian (rabbit) vault particle (PDB: 9Z4W). **f-h** Individual MVP monomers from DbVault (**f**), CpVault (**g**), and rabbit vault (**h**) with C-terminal cap-ring domain excluded.

In contrast to the half-vaults of DbMVP, 2D class averages of CpMVP only showed intact full vaults (CpVault) with 35 monomers making up each vault half. 3D reconstruction with D35 symmetry imposed produced a 2.2 Å resolution reconstruction that measured 640 Å long and 350 Å wide, somewhat shorter and narrower than rabbit vault (**Fig. 2c**). This high-resolution map also allowed us to model much of the protein excluding disordered loops (**Fig. 2d; Extended Data Figure 5b,d; Extended Data Figure 6c**). CpMVP exhibits an MVP structure that consists of 7 repeats of antiparallel β-sheets, an SPFH shoulder domain, and a cap-helix. While the overall length of the cap-helices between CpMVP and mammalian MVP (∼150 a.a.) are similar, CpMVP has two fewer repeat domains, resulting in a reduction of the overall length of CpVault by ∼30 Å (**Fig. 2e, g, h**).

The CpVault cap-helix domain morphology is similar to mammalian vaults, including the presence of an invagination roughly 60 Å away vertically from the end of the shoulder domain. (**Fig. 3a, b**). Interestingly, this invagination is absent in DbVault (**Fig. 3c**); the cap-helix of DbVault instead kinks toward the center at this location, establishing the relatively flattened shape of the DbVault cap-helix domain. The disk-shaped structures at the end of each vault cap (the cap-ring domains) were not resolved in these reconstructions of the entire vault particles due to symmetry mismatch (**Fig. 2a-d**)^6,12^. We will expand on the cap-ring structures in following sections.

**Fig. 3:**
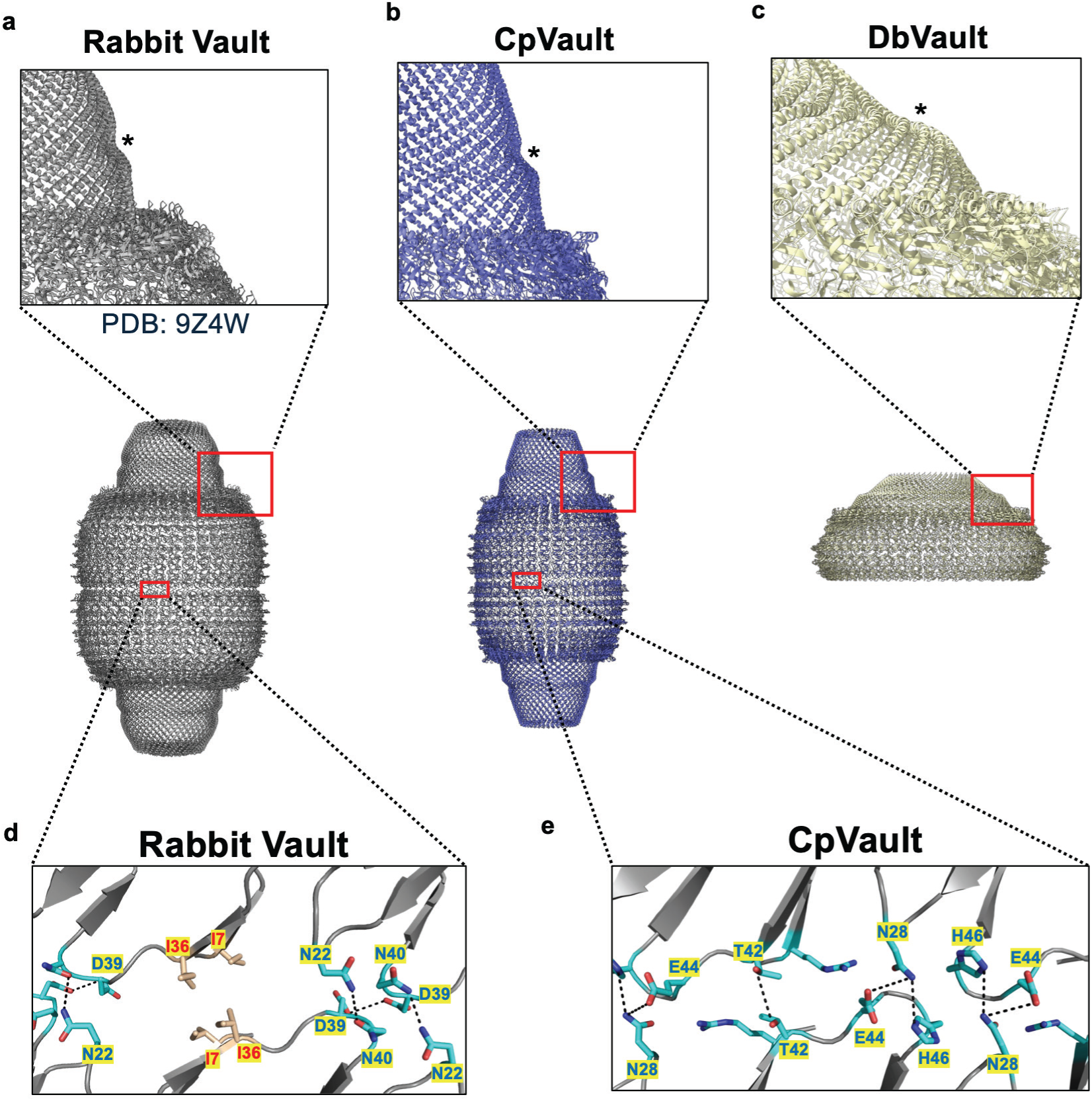
Bacterial vaults exhibit differences in cap-helix structure and waist interactions. **a-c** α-helical kink/invagination within the rabbit vault cap-helix (PDB: 9Z4W) (**a**), DbVault cap-helix (**b**), and CpVault cap-helix (**c**). An asterisk highlights the cap-helix kink of each structure. **d**,**e** Residues and highlighted interactions at the vault waist of rabbit vault (**d**) and CpVault (**e**). Hydrophobic residues are colored wheat and labeled red and polar residues colored cyan and labeled blue. Threshold for selected interactions is 4.5 Å or less.

DbVault and CpVault are held together by interactions between neighboring repeat domains and cap-helices. Inspection of the two repeat domains preceding the shoulder domain of DbVault (R5/R6) and the corresponding region of CpVault (R6/R7) revealed similar, mostly polar interactions between adjacent protomers. In DbVault R5 and R6, residues Y330-E363, Y330-R345, N273-K342, E272-D359, and A271-Y358 apparently maintain close contacts between each chain (**Extended Data Fig. 7a**). In CpVault, similar interactions (N368-R464, R364-D397, Q285-E377, E287-R292, L230-L294, and E229-R295) are present in the corresponding repeats R6 and R7 (**Extended Data Fig. 7b**). At the cap-helix, both CpVault and DbVault MVP protomers are held together by complementary polar and hydrophobic regions that stabilize the cap-helix domains (**Extended Data Fig. 7c, d**).

The rabbit vault waist interface is stabilized by contacts involving both polar and hydrophobic residues (**Fig. 3d**). However, we find that the corresponding interactions in CpVault only involve polar residues, including E44-N28, T42-T42, and N28-H46 (**Fig. 3e**). This contrasts with the network present in the rabbit vault, which utilizes a polar interface between N40-N22 and D39-D39 but also contains hydrophobic residues I7 and I36.

### Symmetry and architecture of the Deltaproteobacteria vault cap

While the structure of the vault barrel/cap-helix has been known for over a decade, the structure of the vault cap-ring domain has been elusive until recently, owing to a symmetry mismatch between the two regions (C39 barrel vs. C13 cap) that complicated cryoEM reconstruction^6,12^. Structures of the human and rabbit vault caps have demonstrated that this 39-fold to 13-fold symmetry transition occurs at the perimeter of the cap-ring domain, which contains α-helices that run perpendicular to the central axis and lead to two concentric β-barrels surrounding a central aperture^5,6,12^.

In our high-resolution reconstructions of DbVault and CpVault described above, the vault cap-ring domains were not resolved, likely due to this same symmetry mismatch^6,12^. To resolve the DbVault cap-ring domain, we performed 3D classification focusing on the cap region of the complex, which revealed 3 evenly distributed classes rotated about the central symmetry axis (**Extended Data Fig. 8a**). Interestingly, each of these classes implied a 16-fold symmetric cap, contrasting the 48-fold symmetry of the DbVault body. Extracting a sub-particle encompassing the cap region and performing further local refinements, we obtained a 2.4 Å resolution map with C16 symmetry imposed (**Extended Data Fig. 8c**). This cap-focused map allowed us to build an average of ∼47 more residues at the C-terminus of the monomer (**Fig. 4a,b; Extended Data Figure 6b**). We also observed tubular densities adjacent to a hydrophobic surface of the DbVault aperture’s cytoplasmic side that we were unable to assign (**Supplementary Fig. 2**).

**Fig. 4:**
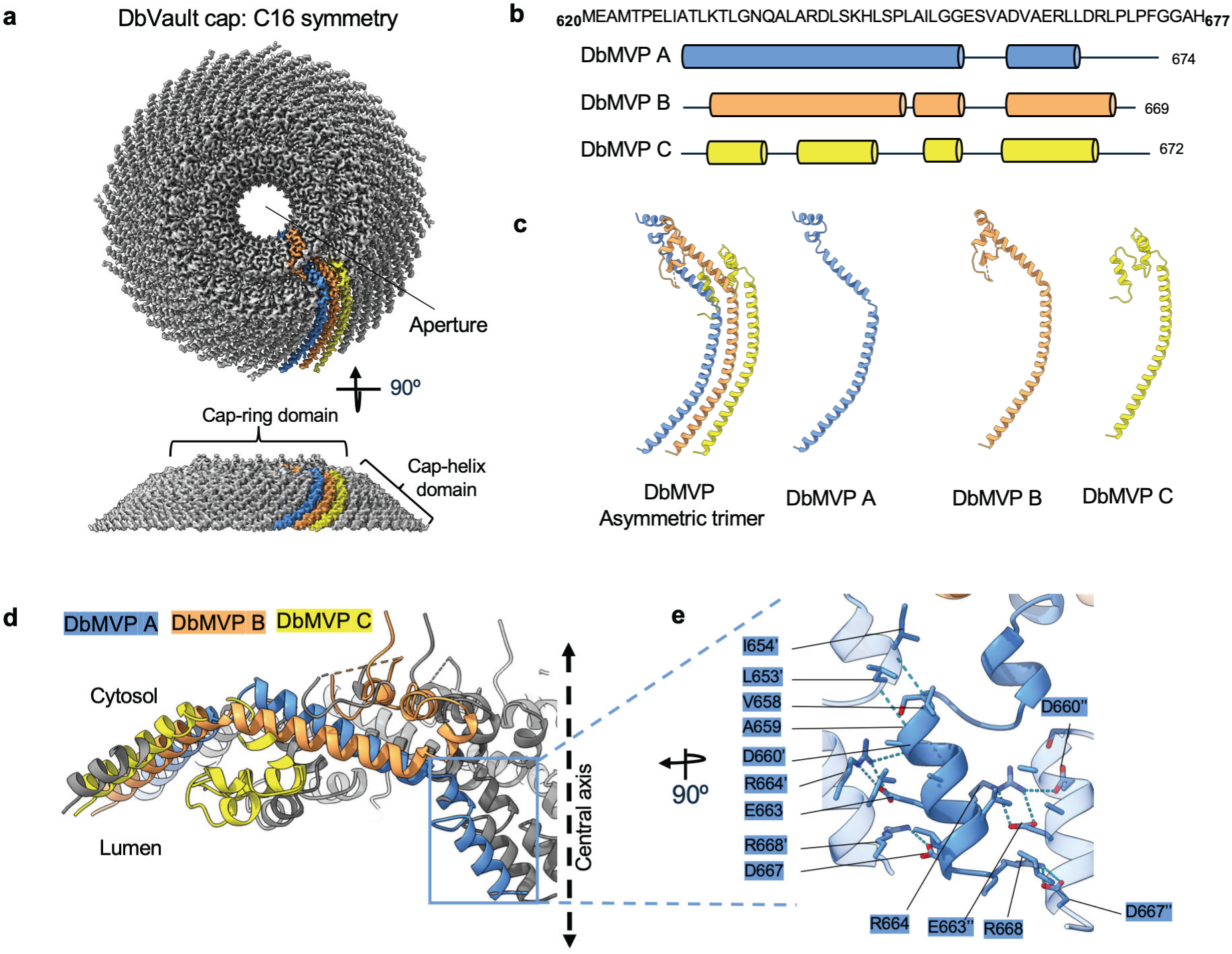
Organization of the DbVault cap. **a** CryoEM reconstruction of the DbVault cap region. The cap-ring and cap-helix domains are labeled. **b** Schematic representation of DbMVP chains that form the cap-ring domain. Cylinders represent α helices. **c** Cartoon representation of the asymmetric trimers that make up the DbVault cap. **d** Sliced view of the DbVault cap-ring domain, viewed perpendicular to the central vault axis which is highlighted as a dotted line. **e** Expanded view of the boxed region in **d**, highlighting unique interactions that stabilize the MVP A helices lining the central aperture. Neighboring MVP A protomers are depicted as semitransparent cartoons and their residues are labeled with apostrophes.

The structure reveals that the 48 protomers comprising each DbVault cap can be broken down into 16 asymmetric trimers. We term the adjacent monomers in the asymmetric trimer MVP A, B, and C and note that the cap-ring domain is made up of exclusively α-helices, in contrast to the α/β structure of mammalian vault cap-ring domains (**Fig. 4b,c**). Structurally, each monomer in the DbVault cap-ring domain assumes a similar α-helical organization until residue A622, where the C48 to C16 symmetry transition occurs. From there, DbMVP C turns toward the interior of the vault, creating a series of four short helices that line the cap interior ‘ceiling’, while helices formed by DbMVP A and B residues 624-647 run roughly parallel with each other toward the central axis (**Fig. 4d**). DbMVP A and DbMVP B diverge after residue 647, where DbMVP A continues toward the center to form two short helices separated by a four-residue linker that line the cytoplasmic side of the vault aperture. DbMVP B turns up toward the vault exterior and folds back in the direction of the cap periphery, making two short helices that settle on top of DbMVP A and a coil that nestles between DbMVP A and DbMVP C of the adjacent asymmetric unit (whose protomers we designate MVP A’, B’, and C’) (**Fig. 4d**).

The DbVault cap-ring domain structure is reinforced by extensive inter- and intra-protomer contacts. For instance, the F763 side chain of DbMVP B remarkably nestles in between three neighboring subunits, making interprotomer contacts with K632 (DbMVP A), E626 and T630 (DbMVP C’), and T633 (DbMVP B’). Salt bridge interactions between neighboring DbMVP B protomers involving R668, E663’ and D660’ propagate across the cap cytosolic surface to stabilize a circular arrangement of short DbMVP B helices ∼45 Å from the central axis (**Extended Data Fig. 9a**). At the DbVault aperture, the C-terminal end of the DbMVP A upper α-helix stacks against the N-terminal end of the lower α-helix of an adjacent DbMVP A subunit. The lower helices are further stabilized across MVP A subunits salt bridges involving E663, R664’, D660’ on one side and R668 and D667’’ on the other side (**Fig. 4e**). We note that DbMVP A and B use sequence-equivalent charged residues (D660, E663, R668) to mediate distinct electrostatic interactions at discrete locations of the cap-ring, highlighting how the symmetry breakdown of the DbVault cap underlies its architecture.

### Symmetry and architecture of the *C. praedator* vault cap

After resolving the symmetry mismatch at the DbVault cap, we sought to apply a similar strategy to the CpVault cap. Initially, we surmised that a different kind of symmetry transition may underlie the CpVault given that the 35 protomers comprising each CpVault cap can’t be evenly divided into trimeric asymmetric units. We hypothesized a hepta- or pentameric asymmetric unit instead could mediate 5- or 7-fold symmetry at the cap. Surprisingly, masked 3D classification of C35-symmetry expanded DbVault particles produced a class displaying 13-fold symmetry in the central region of the cap-ring, like mammalian vaults. Local refinement using a tight mask drawn around cap-ring resulted in a 2.8 Å resolution map in which the innermost region around the central aperture remained less well resolved (**Extended Data Fig. 8b, d**). Another round of local refinement (applying a tighter mask around just the central portion of the cap-ring domain) with C13 symmetry imposed generated a 3.2 Å resolution map with improved connectivity (**Extended Data Fig. 8e**), allowing us to model additional residues in this region (**Fig. 5a, b; Extended Data Fig. 6d**).

**Fig. 5:**
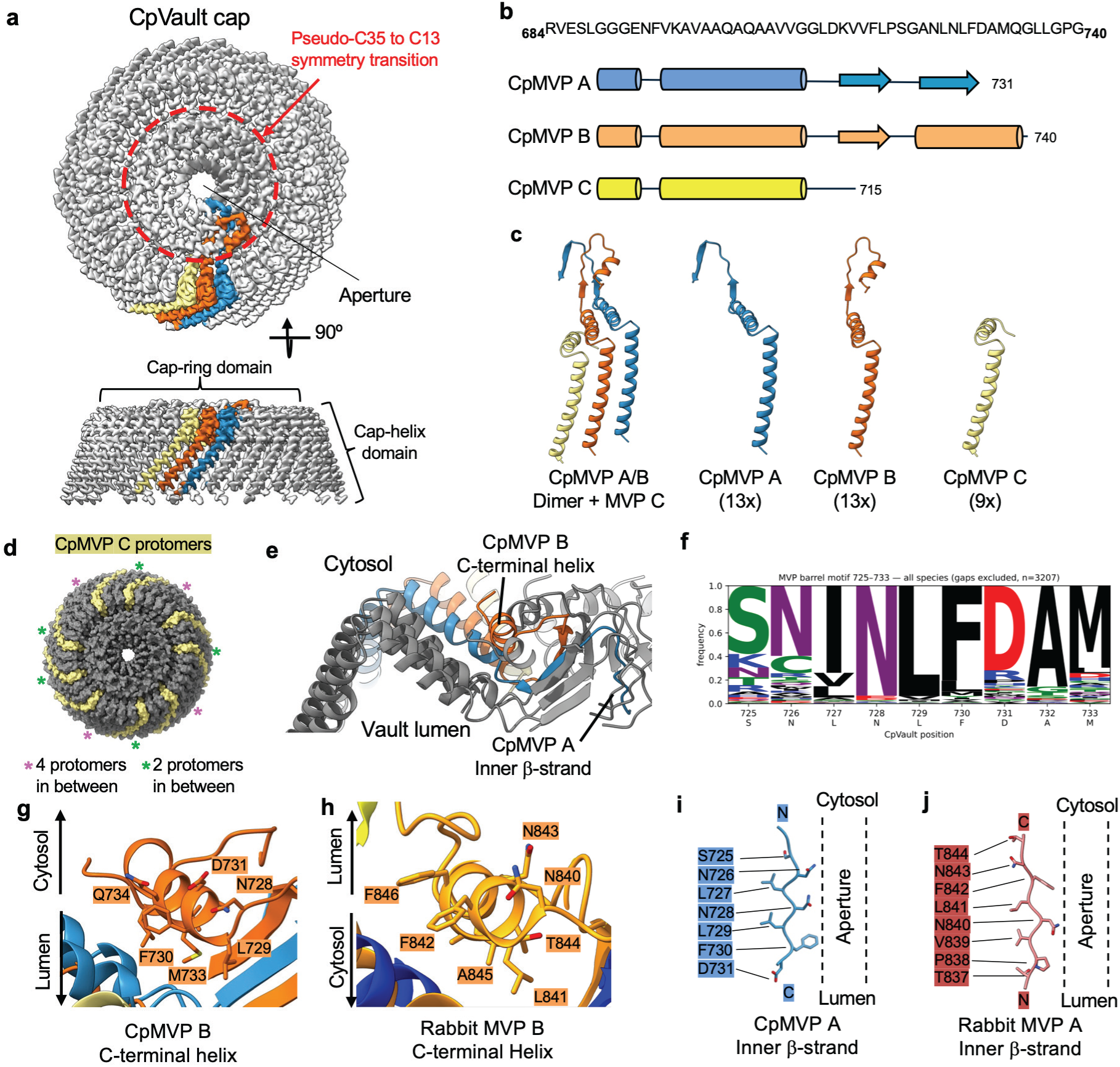
Architecture of CpVault cap and conserved structural role of a C-terminal motif. **a** CryoEM reconstruction of the CpVault cap region. The cap-ring and cap-helix domains are labeled. **b** Schematic representation of CpMVP chains that form the cap-ring domain. Cylinders represent a helices and arrows represent b sheets. **c** Cartoon representations of the asymmetric unit of CpVault cap, which is composed of CpMVP A/B dimers and stochastically excluded CpMVP C. **d** Surface view of the CpVault cap region with CpMVP C protomers colored khaki. Asterisks highlight the varying number of CpMVP A/B protomers slotted between CpMVP C protomers. **e** Cartoon representation of CpVault cap-ring viewed perpendicular to the central axis. **g,h** Expanded views showing conserved C-terminal helices of CpMVP B (**g**) and rabbit MVP B (**h**, PDB: 9Z5N). **i,j** Inner β-strands of CpMVP A and Rabbit MVP A exhibit conserved residues (modeled using DockPrep) pointing toward the aperture.

Analogous to the rabbit vault cap^12^, the CpVault cap is comprised of α-helices on the perimeter that lead to two concentric β-barrel-like structures in the center (**Fig. 5b**; **Extended Data Fig. 6d**). Interestingly, while helices from all 35 protomers contribute to the cap-helix domain, the central 13-fold symmetric portion of the cap-ring is composed of 26 protomers, 13 of which participate in the inner β-barrel. Using a similar nomenclature to DbVault, we designate the 13 protomers that make up the inner β-barrel as CpMVP A, the remaining 13 protomers that contribute to the outer β-barrel as CpMVP B, and the 9 protomers excluded from the cap-ring center as CpMVP C (**Fig. 5c**). The uneven distribution of protomers results in a cap-ring domain with 9 CpMVP C protomers slotted in between 2 or 4 CpMVP A/B protomers in a seemingly stochastic fashion (**Fig. 5d**), in contrast to the regularly arranged asymmetric trimers that make up the DbVault and mammalian vault caps. The symmetry transition from pseudo-C35 to C13 occurs at V712, after which CpMVP C drops into the vault lumen and loses cryoEM density, presumably due to disorder. CpMVP A and CpMVP B assemble into an outer β-barrel starting at E716 before separating at P722, where CpMVP B turns away from the center and into a short α-helix that rests on the outer surface of the vault cap-ring, and CpMVP A loops back toward the central axis where it forms an inner β-barrel aperture similar to what has been observed in mammalian vault caps^6,12^ (**Fig. 5e**). This contrasts with the DbVault cap where we observe α-helices lining the central aperture.

### Architectural principles of cap domains in vaults and other SPFH-type complexes

We noticed that the asymmetric folding of CpMVP C-terminal residues (S725-Q734) into an inner β-strand (in CpMVP A) and short α-helix (in CpMVP B) (**Fig. 5e**) was strikingly similar to the conformations of the rabbit MVP A/B C-termini^12^, suggesting this feature could be broadly conserved. Indeed, sequence analysis of this region revealed that a C-terminal sequence motif centered at CpMVP residue L729 (ϕ, N, L, F, ζ, A; where ϕ is any hydrophobic residue and ζ is any charged residue) is conserved in 61.9% of MVP homologs (**Fig. 5f, Extended Data Fig. 10a**). In CpMVP B, residues N728, L729, and F730 (N, L, F in the motif) reside at the start of the C-terminal helix, with L729 pointing toward the outer β-barrel (**Fig. 5g, Extended Data Fig. 9b**). Conserved residues in rabbit MVP B (N840, L841, F842) adopt corresponding positions in its C-terminal helix, including the projection of L841 toward the outer β-barrel of the rabbit vault cap-ring (**Fig. 5h**). The inner β-strand of CpMVP A (composed of residues S725-D731) had weaker cryoEM density, consistent with the rabbit vault cap inner β barrel density^12^. We used DockPrep in ChimeraX^49–51^ to model likely side chain rotamers for amino acids of the inner CpMVP A β-strand. The side chains of N728 and F730 (of the N, L, F motif) were modeled to point into the CpVault aperture (**Fig. 5i**) and form a ∼14 Å constriction between opposing subunits. Remarkably, the corresponding residues in rabbit MVP A (N840 and F842) also point toward the central aperture (**Fig. 5j**) to generate a pore constriction of roughly the same width as CpVault (**Fig. 6a,b**).

**Fig. 6:**
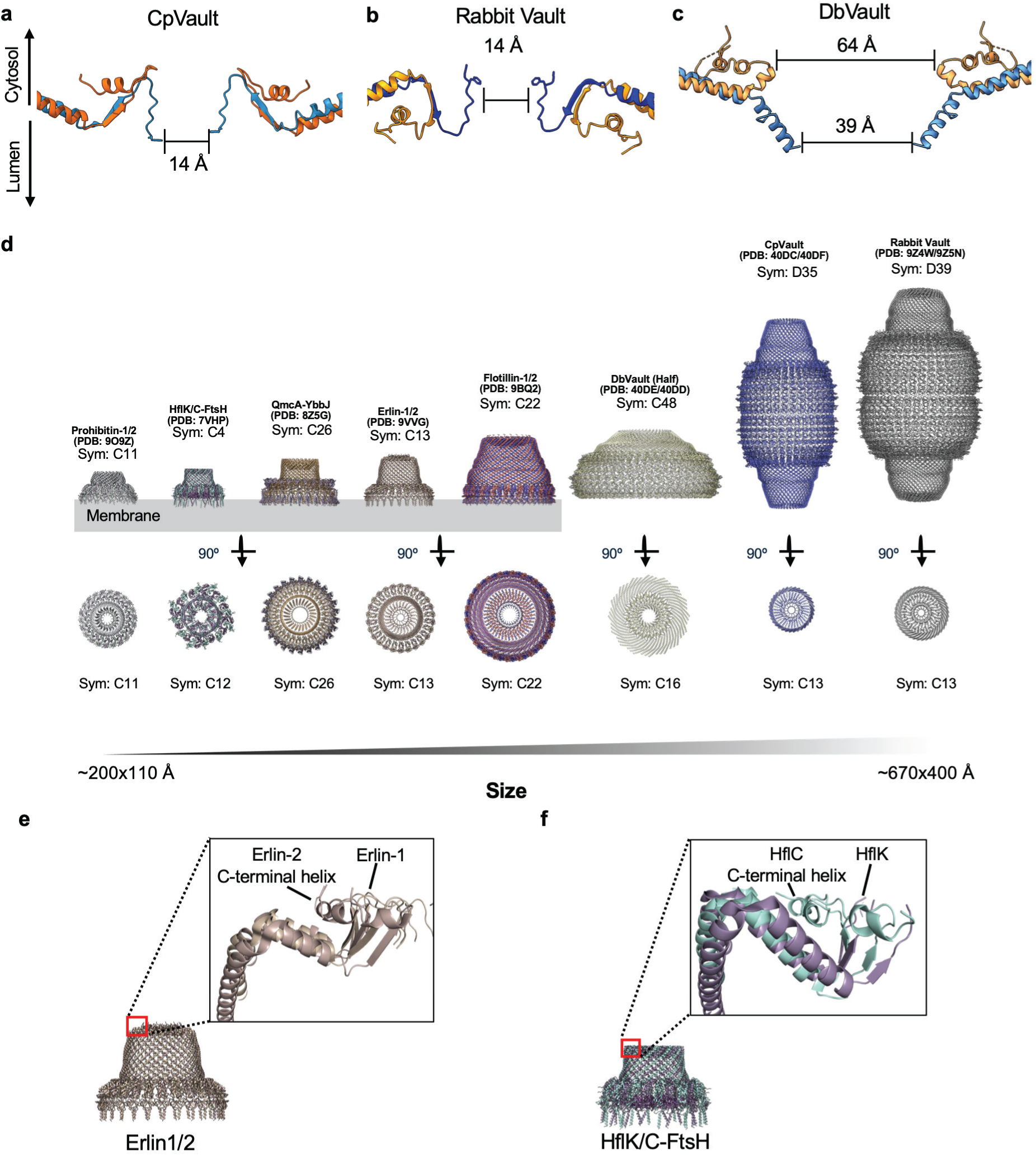
Structural features of vaults and SPFH protein complexes. **a-c** Structural comparison of the CpVault (**a**), rabbit Vault (**b**) and DbVault (**c**) central apertures. Opposing MVP A and B monomers are shown for clarity. Aperture widths between conserved phenylalanine residues are shown in **a** and **b**. Minimum aperture widths between opposing MVP A and B subunits are shown in **c. d** Comparisons of membrane-bound SPFH family members: Prohibitin-1/2 complex (11 Prohibitin-1 + 11 Prohibitin-2, PDB: 9O9Z), HflK/C-FtSH complex (12 Hflk + 12 HflC + 24 FtsH, PDB: 7VHP), QmcA-YbbJ complex (26 QmcA + 26 YbbJ, PDB: 8Z5G), Erlin-1/2 complex (13 Erlin-1 + 13 Erlin-2, PDB: 9VVG), Flotillin-1/2 complex (22 Flotillin-1 + 22 Flotillin-2, PDB: 9BQ2), and cytosolic SPFH members: DbVault (PDB: XXX), CpVault (PDB: XXX), and rabbit vault (PDB: 9Z4W). All proteins are to scale. Figure partially created with biorender.com. **e-f** Structural views of the caps of the Erlin1/2 complex (**e**) and HflK/C-FtsH (**f**). Expanded views highlight topologies of the C-terminal regions.

Our observation that the N, L, F-containing C-terminal motif doubles as an α-helix (in MVP B) and inner β-strand (in MVP A) in both CpVault and mammalian vault suggests it may underlie conserved cap-ring architectures across many MVP/vault homologs. However, despite the similarities in fold, the C-terminal topologies of MVP A/B are inverted in CpVault (MVP A in vault lumen and MVP B cytosolic) and rabbit vault (MVP A cytosolic and MVP B in vault lumen) (**Fig. 5g-j, 6a-b**). This results in opposite surfaces of the cap-ring facing the interior and exterior of CpVault and rabbit Vault. It is also notable that this C-terminal motif is absent in DbMVP (**Extended Data Fig. 10b**), consistent with its divergent cap-ring symmetry and structure. The central α-helical aperture of the DbVault cap-ring measures ∼64 Å between DbMVP B subunits at the cytosolic mouth and ∼36 Å between DbMVP A subunits at the luminal side of the aperture (**Fig. 6c**), significantly wider than CpVault and rabbit vault. The diverse architectures, topologies, and aperture widths of bacterial and mammalian vault cap-rings may have implications for the nature of molecules that can interact with the cap or pass through its aperture.

While other SPFH-type complexes lack repeat domains, their caps share structural similarities with vault caps (**Fig. 6d**)^12,13,52–55^. For example, Erlin-1/2 and HflK/C-FtSH are each composed of two distinct protomers that form a cap-ring domain outer β-barrel, analogous to CpMVP A and B of CpVault. Beyond this β-barrel, both the Erlin-2 subunit of the Erlin-1/2 complex and HflC of HflK/C-FtSH complex form C-terminal helices that packs against the cytosolic side of the cap in a highly similar manner to the CpMVP B C-terminus (**Fig. 6e,f**). The Erlin-1 subunit continues toward the central axis of the complex to form a possible inner β-barrel analogous to CpMVP A, while the HflK C-terminus is unresolved (**Fig. 6e,f**). Notably, the cap-rings of Flotillin-1/2 and Prohibitin-1/2 follow an equivalent topology involving an MVP B-like subunit with its C-terminus on the complex exterior, akin to CpVault and DbVault but opposite to mammalian vaults. The similarities between the caps of bacterial vaults and SPFH family complexes may reflect a retention of topological features from an ancestral SPFH-like assembly.

## Discussion

Through phylogenetic, biochemical, and structural analyses, we identify bacterial MVP homologs that form vault or vault-like structures. Our phylogenetic analysis shows that a small subset of bacterial genomes, mostly within the Cyanophyceae and Myxococcia classes, encode MVP homologs that vary in sequence length and predicted repeat domain number. We successfully expressed and purified six bacterial MVP homologs, two of which resulted in high resolution cryoEM structures of vault complexes (DbVault and CpVault). Three other MVP homologs (MbMVP, CaMVP, PpMVP) formed higher-order vault-like assemblies that appeared to be fractured, possibly because our expression/purification conditions did not provide the optimal environment or host factors required for intact vault complex isolation. Though some species have been reported to have multiple MVP paralogs that may be needed to properly form the vault complex^37–40^, we were not able to identify paralogs of the MVPs experimentally screened in this study. Nonetheless, our observations on a subset of homologs suggest that bacterial MVPs have an overall propensity to form vault-shaped assemblies.

Both DbVault and CpVault exhibit structural divergence from mammalian vaults, with the former being more distinct. It is unclear why DbMVP only assembled into half-vaults. While mammalian vaults are known to participate in half-vault exchange^16^, intact vaults are the predominant form found in solution^6,11^ and in cells^38,56^. Furthermore, clear homology between R1 domains of DbVault (res. 1-91) and rabbit vault (res. 1-50) (**Extended Data Fig. 10b**) indicates the DbVault “waist” is amenable to full vault formation, making it difficult to conclude that the half-vault is the biologically relevant form. In addition to its divergent half-vault morphology, our DbVault structure displays several unique features: 48/16-fold symmetries of the body/cap, a flattened cap-helix domain architecture, and exclusively α-helical central cap-ring structure (**Figs. 2, 3, 4, 6**).

CpVault forms an intact vault structure with 35-fold symmetry at the barrel that breaks into 13-fold symmetry at the cap-ring domain (**Fig. 5a-d**). The CpVault cap contains four fewer CpMVP C protomers than A and B, resulting in an irregular arrangement of 13 CpMVP A/B dimers and 9 CpMVP C protomers that contrasts the evenly distributed asymmetric homotrimers observed in DbVault and mammalian vault caps. This demonstrates for the first time that the vault cap-ring symmetry does not have to be a factor of the vault body symmetry, suggesting that vault assembly can tolerate a wider range of symmetries than may have been previously assumed. We found that a well-conserved C-terminal motif (ϕ, N, L, F, ζ, A) underlies the dual α/β fold surrounding the central cap aperture of CpVault and mammalian vault, though their topologies are inverted (**Fig. 5e-h**). We speculate that MVP homologs harboring this C-terminal motif (which makes up 61.9% of homologs identified in this study) may all share similar C13-symmetric cap-ring folds. This motif is not conserved in DbVault, which has a wider α-helical cap-ring aperture with C16 symmetry. Vault cap-rings with diverse topologies and architectures may underlie different functions across species, including dictating diffusion through the central aperture as has been proposed previously^6^.

Our DbVault and CpVault structures consist of MVP subunits with fewer than the canonical 9 repeat domains encoded by mammalian MVPs. To probe whether specific repeat domains were under selective pressure to be excluded, we compared the sequence identity of each human repeat domain (based on Uniprot-annotated domains) to that of our entire data set of MVP homologs and to bacterial sequences only (**Extended Data Fig. 10c, d**). We find that sequences corresponding to human R3-R5 are often absent in bacteria, suggesting that loss of these regions underlies the decreased length of certain bacterial MVP homologs. Indeed, sequence alignments of rabbit MVP against CpMVP, MbMVP, and CaMVP show major gaps at portions corresponding to R4 and R5 (**Extended Data Fig. 10a; Supplemental Fig. 3**). On the other hand, a sequence corresponding to R5 is absent in DbMVP, and partial sequences of R2/R3, and R4/R6, each appear to constitute a single repeat domain (**Extended Data Fig. 10b**). This highlights the possibility that prokaryotic MVPs may have evolved from an ancient ancestor and selectively lost specific repeat domains over time. Interestingly, we note that vPARP binds to the interior of human vault via contacts contributed mostly from R4^3^. Since most prokaryotes lack a PARP homolog^57^, we speculate that selective loss of the R4 domain could have been a mechanism utilized by bacteria to conserve resources while also maintaining a functional, albeit smaller, vault complex. Ultimately, *in situ* analyses of these complexes within their native cellular environment are critical to further define the native architecture of bacterial vaults and uncover possible biological functions.

Our bioinformatics analyses identified additional divergent candidate MVPs that may be of interest for future study. These include many eukaryotic MVP sequences shorter than typical mammalian MVPs (27.1% of eukaryotic sequences were in the 580-850 a.a. range) as well as 22 archaeal MVP homologs^58^. Elucidating these MVP assemblies will expand our knowledge of vault diversity well beyond the bacterial structures we have reported here.

## Methods

### Phylogenetic analysis of MVP homologs

MVP homologs were identified by combined sequence- and structure-based searches. A conserved 36-aa peptide query was first derived from human MVP (NP_005106.2): the NCBI Nucleotide database^59^ was searched for “major vault protein” (records 1,000 bp–10 Mb), and the 4,023 hits were filtered to 3,352 sequences sharing >30% identity and coverage with human MVP, aligned with MUSCLE^60^, and scored for per-column conservation (custom pipeline) to select the most conserved window. Two BLASTp searches (BLAST+ v2.12.0^61^) against the NCBI non-redundant protein database were performed using full-length human MVP (NP_005106, 893 aa) and a conserved MVP peptide as queries, yielding 3,511 unique accessions. In parallel, two Foldseek^47^ searches against the AlphaFold Database used the human MVP structure (AF-Q14764) and a prokaryotic MVP structure (AF-A0A3A8K409) as queries; hits were retained at ≥25% sequence identity and E-value <1×10⁻²⁰, giving 1,700 candidates. The two sets were reconciled at the sequence level using DIAMOND^62^ (v0.9.14) against a database built from the BLAST hits, with Foldseek candidates matching a BLAST sequence at ≥99% identity considered redundant. The merged, non-redundant dataset comprised 3,959 MVP homologs. Taxonomic lineages were obtained from NCBI Taxonomy (BLAST hits) and UniProt (Foldseek-only candidates) and harmonized against the NCBI lineage vocabulary.

Taxonomic distribution was visualized as a cladogram-style tree built from the NCBI/UniProt-harmonized lineages (domain through class), drawn in R. The cladogram (Fig. 1a) displays the 3,825 of 3,959 homologs whose lineages were fully resolved from domain to class from NCBI; the 134 sequences lacking a class-level assignment were excluded from the cladogram. Of the sequences displayed, 298 were prokaryotic (290 Bacteria, 8 Archaea). For protein length distributions, fused sequence from misannotated gene models was removed from the full-length proteins by a symmetric BLAST-anchor de-fusion: the human cap (aa 521–893) anchored removal of C-terminal fusions, and full-length human MVP (aa 1–893) anchored removal of N-terminal fusions, in each case trimming beyond the inferred MVP boundary plus a 60-aa margin while preserving tandemly duplicated MVPs. The resulting lengths were plotted by taxonomic group in R.

The conserved C-terminal cap domain was used as the phylogenetic substrate. Caps were extracted by an anchor-based approach: the human R9 repeat (aa 457–520) was used as a BLASTp query against each homolog and the sequence downstream of the R9 match was taken as the cap; sequences lacking an R9 match were rescued using the human cap (aa 521–893) as a probe. C-terminally fused sequence was removed using the cap probe as a symmetric C-terminal anchor. Caps were filtered by length (165–900 aa), retaining 3,331 sequences including 288 bacterial and 21 archaeal caps. The sequences were aligned with MAFFT v7.490^63^ (--auto) and trimmed of gap-rich columns with trimAl v1.4^64^. A maximum-likelihood tree was inferred with IQ-TREE v2.1.4^65^ under the site-heterogeneous LG+C20+F+G mixture model using the posterior mean site frequency (PMSF) approximation with a FastTree v2.1.10^66^ (LG+G) guide tree; branch support was assessed with 1,000 bootstrap replicates and the SH-aLRT test. The tree was rooted using the minimum-variance criterion (MinVar-Rooting)^67^, which is robust to among-lineage rate variation and long-branch effects. Branch lengths in the circular phylogram were square-root–transformed for display.

Sequence identity of selected MVPs against human MVP are determined by alignment with R Biostrings/pwalign package (pairwiseAlignment, type=“global”) using the BLOSUM62 substitution matrix with gap penalties of opening = 10 and extension = 0.5. Sequence logo for CpMVP position 725-733 was generated by aligning all 3,959 MVP homologs with MAFFT v7.490 (--auto) using NCBI accession WP_120582404 as reference. A probability logo was produced by logomaker, excluding gaps and normalizing each column to 100%. Sequence alignment between rabbit MVP and DbMVP, CpMVP, CaMVP, or MbMVP in Extended Data Fig. 10 and Supplementary Fig. 3 was performed using MacVector (version 18.7.3, Macvector, Inc.)

### Expression and purification of MVP homologs

MVP homologs were synthesized and cloned into pET29(a) vectors containing either a C-terminal tobacco etch protease cleavage site (TEV) followed by a StrepII tag (CpMVP, CaMVP, OsMVP, PpMVP, MbMVP) or a hexahistidine tag fused directly to the C-terminus (DbMVP). Plasmids were transformed into BL21 (DE3) *E. coli* and grown in a 20 mL LB culture overnight containing 50 μg/mL of kanamycin. 5mL of overnight culture was utilized to inoculate a 2L flask containing 500 mL of LB media supplemented with 50 μg/mL of kanamycin and was incubated at 37°C and 225 rpm shaking until OD reached at least 0.6. The shaker temperature was then reduced to either 22°C (DbMVP, MbMVP, and PpMVP) or 18°C (CaMVP, CpMVP, and OsMVP) and after 20 minutes, IPTG was added to either a 1mM (DbMVP, MbMVP, and PpMVP) or 300 μM (CaMVP, CpMVP, and OsMVP) final concentration. Cultures were incubated for 16-24 hours before being harvested by centrifugation at 10,000 x g for 10 minutes. Cell pellets were frozen at -80°C until purification.

The compositions of buffers used for MVP purification are listed in **Supplementary Table 1**. Cell pellets were thawed and resuspended in 3 mL of lysis buffer per gram of wet pellet and sonicated using a Banson Digital Sonifier 450 device equipped with a microtip at 15% amplitude with pulses of 15 s on 15 s off, for a total of 3 minutes of sonic disruption. The lysate was then centrifuged at 9660 x g for at least an hour. DbMVP lysate was loaded onto a 60mL HisPure Cobalt gravity column that was pre-equilibrated with wash buffer. The column was washed with 5 bed volumes of wash buffer and eluted with 2 bed volumes of elution buffer. Eluent was concentrated with a 100kDa MWCO concentrator (Amicon). DbMVP protein was stored at -80°C in elution buffer. Protein concentration was determined by A280 using a generalized extinction coefficient of 1 Abs = 1 mg/mL.

For all other constructs, the resulting supernatant was passed over a 5mL StrepTrapXT column at a flow rate of either 5 mL/min (MbMVP) or 0.5 mL/min (PpMVP, CaMVP, CpMVP, OsMVP) using an Akta Pure or Akta Avant purification system. The column was washed with 20 CV with wash buffer at a flow rate of 5mL/min and eluted with 5 CV of elution buffer at a flow rate of either 5mL/min (MbMVP) or 1mL/min (PpMVP, CaMVP, CpMVP, OsMVP). The resulting fractions with high A280 absorbance were pooled and concentrated with a 100 kDa MWCO concentrator (Amicon) and subsequently stored at -80°C. Protein concentration was determined by A280 using a generalized extinction coefficient of 1 Abs = 1 mg/mL.

After affinity chromatography, CaMVP and OsMVP were further purified by a discontinuous sucrose gradient. The discontinuous sucrose gradient was made by mixing sucrose with MQ water (v/v) to yield 60%, 50%, 45%, 40%, 30% and 20% sucrose solutions. MVP from the affinity step was gently overlayed on top of the gradient, and samples were centrifuged in a fixed angle rotor (Type 45 Ti) at 78,000 x g for 16-20 hours. CaMVP exhibited particles that equilibrated at the 60% sucrose layer which was pooled for further screening. OsMVP exhibited particles that equilibrated between 20-40% sucrose, and the 40% fraction was pooled for further screening. Fractions were concentrated and exchanged into 50mM Tris HCl pH 7.5, 75mM NaCl, 0.5 MgCl_2_ using a 100kDa MWCO concentrator (Amicon).

### CryoEM sample preparation and data collection

The DbMVP sample was exchanged into PBS while all other MVP samples remained in their respective elution buffers. The DbMVP sample was prepared at 2.6 mg/ml while the CpMVP sample was prepared at 8.1 mg/ml. CFlat 2/1 grids were plasma cleaned in a Solarus II (Gatan) using a H2/O2 gas mixture. 0.01% fluorinated octyl-maltoside (FOM) was added to the concentrated samples to mitigate preferred orientation. 3 µl of sample solution was added to the CFlat grids before plunge freezing (blot time 4 seconds, blot force -1) into liquid ethane cooled by liquid nitrogen in a Vitrobot Mark IV operated at 4°C and 100% humidity.

Grids were loaded into a Titan Krios G3i electron microscope equipped with a BioQuantum K3. Images were collected in counting mode at either a magnification of 105kx, calibrated pixel size of 0.839 Å (DbMVP), or 81kx, calibrated pixel size of 1.061 Å (all other samples). For DbMVP data collection, a defocus range of −0.8 to −2.2 µm was used, the energy filter was inserted with a width of 20 eV, and the 100 µm objective aperture was inserted. Each movie was dose fractionated into 46 frames over a 2.0 s exposure and had a total dose of ∼44 e-/Å2. For samples collected at 81kx, a defocus range of −0.8 to −2.2 µm was used, the energy filter was inserted with a width of 25 eV, and the 100 µm objective aperture was inserted. Each movie was dose fractionated into 46 frames over a 2.0 s exposure and had a total dose of ∼41 e-/Å2. Datasets were preprocessed in a similar manner. Briefly, DbMVP movies were preprocessed in CryoSPARC Live and the patch motion corrected micrographs were imported into CryoSPARC^68^ for Patch CTF estimation. All other samples were processed without CryoSPARC Live. Micrographs containing thick ice and poor CTF resolution estimations (>7Å for DbMVP and >5Å for all other datasets).

### CryoEM data processing of DbVault

A random 750 micrograph subset of the larger dataset was used for blob picking (circular blob, 350-450Å), from which 2D templates were generated via iterative 2D classification. These templates were then used for template picking for the whole dataset. 600K particles were extracted (box size 1024, cropped to 64 pix) then curated using iterative 2D classification and ab initio refinement (K=3, class similarity = 0) yielding 120K particles. This set of particles was then extracted using a box size of 1024, cropped to 512 px, and refined in C1 using non-uniform refinement yielding a 4.2 Å map. These particles were then used to train a topaz^69^ model using a 750-micrograph subset. Topaz picked particles were curated as described above, the two particle stacks were combined, and duplicates were removed using a minimum separation distance of 100 Å yielding 130K particles. Particles were extracted at a box size of 1024 binned to 256 for subsequent 2D classification and ab initio classification which yielded 135K particles. Particles were then extracted at a box size of 1024 binned to 672, aligned to C48 symmetry, and refined to 2.89 Å with C48 symmetry imposed in non-uniform refinement. Heterogenous refinement with 2 identical DbMVP maps and one noise map resulted in a stack of 95K particles that were subjected to CTF refinement, reference-based motion correction (RBMC), Ewald sphere correction (EWS), and local refinement, further improving the resolution to 2.79 Å.

The cap region of the DbVault had smeared densities, suggesting pseudosymmetry. 3D classification without alignment (3 classes, 6Å filter resolution) focused on the vault region revealed 3 equal classes with 16 protomer helices pointing toward the vault lumen rotated about the Z-axis. Aligning all particles to one of these maps and performing local refinement with C16 symmetry imposed yielded a 2.6 Å resolution map with clear side chain densities for the previously unresolved vault cap. Removal of particles with poor per-particle scale factors reduced the particle stack size to 85K. Local refinement, with C16 symmetry imposed, of this particle stack yielded a 2.42 Å resolution map that was used for model building.

### CryoEM data processing of CpVault

To obtain a global reconstruction of the CpMVP sample, a similar schema to DbMVP was followed, with a larger blob for the initial template generation step (circular blob, 350-450Å). Curated template picked particles yielded a 3.1 Å resolution D35 symmetric map (box size 1024, not binned). Training a topaz model using this particle set followed by extensive curation yielded a particle stack that contained 219K particles. CTF refinement, EWS correction, RBMC, and homogenous refinement of this stack generated from a 2.2 Å resolution D35 symmetric map (box size 1024, not binned).

Initial attempts to resolve the asymmetry at the CpVault cap using 3D classification (K=2-100, filter resolution 3-12 Å) failed to produce classes with improved density. To overcome this, we isolated a subparticle focused on just one cap region and performed C35 symmetry expansion yielding 7.6 M particles. 3D classification of the symmetry expanded particle set (K=7, filter resolution 3 Å) yielded one class (1.7 M particles) that displayed 13 helical densities pointing into the vault interior. C1 local refinement yielded a 2.64 Å resolution map, although the aperture region of the cap remained less well resolved. 3D classification using an even tighter mask focused only on the region in the vicinity of the central aperture produced five classes rotated around the central axis containing 1.3M particles. Aligning the five 3D volumes and removing duplicate particle positions resulted in a stack of 131K particles. After local and global CTF refinements, local refinement of this particle stack with C13 symmetry imposed yielded a 3.24 Å resolution map with improved density for the aperture region. A composite map generated in Phenix using the C1 and the C13 refined maps was used for building the cap^70^.

### Model building and refinement

For model building all maps were sharpened using Phenix Autosharpen, supplying the unsharpened maps output from CryoSPARC. Models were iteratively built manually in COOT 0.9.8.94 and 1.1.19 ^71^ and real space refined in PHENIX 1.21.166^70^. The model of the DbVault “barrel” was built into the C48-symmetric map of the DbVault particle using an AlphaFold prediction of the monomer from https://alphafold.ebi.ac.uk/ (Uniprot: A0A661NLQ3) as an initial model^44–46^. This model was adjusted to include residues 26 to 311 (R1 – R5), 322-465 (R6 – partial shoulder), and 475-621 (partial shoulder – cap-helix). Sidechains of R1 and R2 (residues 26-123) were truncated due to weak density. An intrinsically disordered internal loop is also present from 312-321, which connects the two terminal repeats before the shoulder domain—R5 and R6. Notably, this loop corresponds to an unmodeled loop that connects R8 and R9 in mammalian vaults^3–7,9,10,12^. The C16-symmetric locally refined map of the DpVault cap enabled *de novo* modeling of residues 601-669 of MVP A, 601-674 and 687-698 of MVP B, and 601-673 of MVP C.

The structure of CpVault was built using a top scoring prediction from AlphaFold2 Monomer as an initial model^44^. The D35-symmetric map of the CpVault “barrel” was cropped using ChimeraX to reduce computation cost then used to build the repeat domains, shoulder, and cap helix (residues 12-332 and 355-708). A composite map generated in phenix^70^ from the C1 and C13 locally refined cap-ring maps was used to for *de novo* modeling of cap-ring residues 654-731 of MVP A, 654-740 of MVP B, and 654-713 of MVP C. All structural figures were generated in either ChimeraX^49–51^ or PyMol^72^. To model the unresolved rotamers of the inner β-barrel in the CpVault, DockPrep was utilized in ChimeraX. Rotamers were modeled based on Dunbrack library73. CryoEM structure geometry validation statistics were calculated using MolProbity as implemented in PHENIX^74^. CryoEM collection and refinement statistics are shown in Table 1.

**Table 1.** Cryo-EM data collection, refinement and validation statistics.

|  | DbVault<br>(EMDB-XXXX)<br>(PDB XXXX) | DbVault Cap<br>(EMDB-XXXX)<br>(PDB XXXX) | CpVault<br>(EMDB-XXXX)<br>(PDB XXXX) | CpVault Cap<br>(EMDB-XXXX)<br>(PDB XXXX) |
|---|---|---|---|---|
| **Data collection and processing** |  |  |  |  |
| Magnification | 105kx | 105kx | 81kx | 81kx |
| Voltage (kV) | 300 | 300 | 300 | 300 |
| Electron exposure (e–/Å<sup>2</sup>) | 40 | 40 | 42 | 42 |
| Defocus range (μm) | -0.8 to -2.2 | -0.8 to -2.2 | -0.8 to -2.2 | -0.8 to -2.2 |
| Pixel size (Å) | 0.839* | 0.839* | 1.0641* | 1.0641* |
| Symmetry imposed | C48 | C16 | D35 | C1 and C13** |
| Initial particle images (no.) | 600k (blob) / 144k (Topaz) |  | 1.05M (blob) / 299k (Topaz) |  |
| Final particle images (no.) | 95k | 85k | 219k | 1.7M (C1) / 131k (C13) |
| Map resolution (Å) | 2.8 | 2.4 | 2.2 | 2.6 (C1) / 3.2 (C13) |
| FSC threshold | 0.143 | 0.143 | 0.143 | 0.143 |
| **Refinement** |  |  |  |  |
| Initial model used | AlphaFold (A0A661NLQ3) | de novo | AlphaFold2 Prediction | de novo |
| Map sharpening *B* factor (Å<sup>2</sup>) | 82.04 | 79.55 | 47.14 | 74.80 (C1) / 149.23 (C13) |
| Model composition |  |  |  |  |
| Non-hydrogen atoms | 200976 | 27328 | 375200 | 20480 |
| Protein residues | 27264 | 3648 | 47250 | 2685 |
| Ligands | N/A | N/A | N/A | N/A |
| *B* factors (Å<sup>2</sup>) |  |  |  |  |
| Protein (min/max/mean) | 13.00/205.43/102.25 | 19.01/120.35/55.01 | 35.42/162.33/77.41 | 12.33/198.85/94.73 |
| Ligand | N/A | N/A | N/A | N/A |
| R.m.s. deviations (# > 4σ) |  |  |  |  |
| Bond lengths (Å) | 0.004 (0) | 0.004 (0) | 0.005 (0) | 0.001 (0) |
| Bond angles (°) | 0.900 (0) | 0.562 (1) | 0.988 (11) | 0.299 (1) |
| Validation |  |  |  |  |
| MolProbity score | 1.24 | 1.56 | 1.43 | 1.02 |
| Clashscore | 2.44 | 4.52 | 7.96 | 2.42 |
| Poor rotamers (%) | 0.63 | 1.94 | 0.88 | 0.63 |
| Ramachandran plot |  |  |  |  |
| Favored (%) | 96.64 | 97.44 | 98.66 | 99.04 |
| Allowed (%) | 3.36 | 2.56 | 1.19 | 0.96 |
| Disallowed (%) | 0 | 0 | 0.15 | 0 |

\*For the CpVault dataset, all 3D maps were refined at the calibrated pixel size of 1.0641Å/pix. For the DbVault dataset, the cap was refined into a map reconstructed from unbinned particles (0.839 Å/pix) while the DbVault body was refined into a map reconstructed from particles binned to 1.2785 Å/pix.
\*\*See methods for how symmetry was imposed to determine the final map.

## Data Availability

All data needed to evaluate the conclusions in the paper are present in the paper, the supplementary materials, or have been deposited in the PDB and EMDB (accession codes: XXXX, XXXX, XXXX, and XXXX.) Regeneron materials described here may be made available to qualified, academic, non-commercial researchers through a material transfer agreement upon request at https://regeneron.envisionpharma.com/vt_regeneron/. For questions about how Regeneron shares materials, contact by.

---

**Extended Data Fig. 1:**
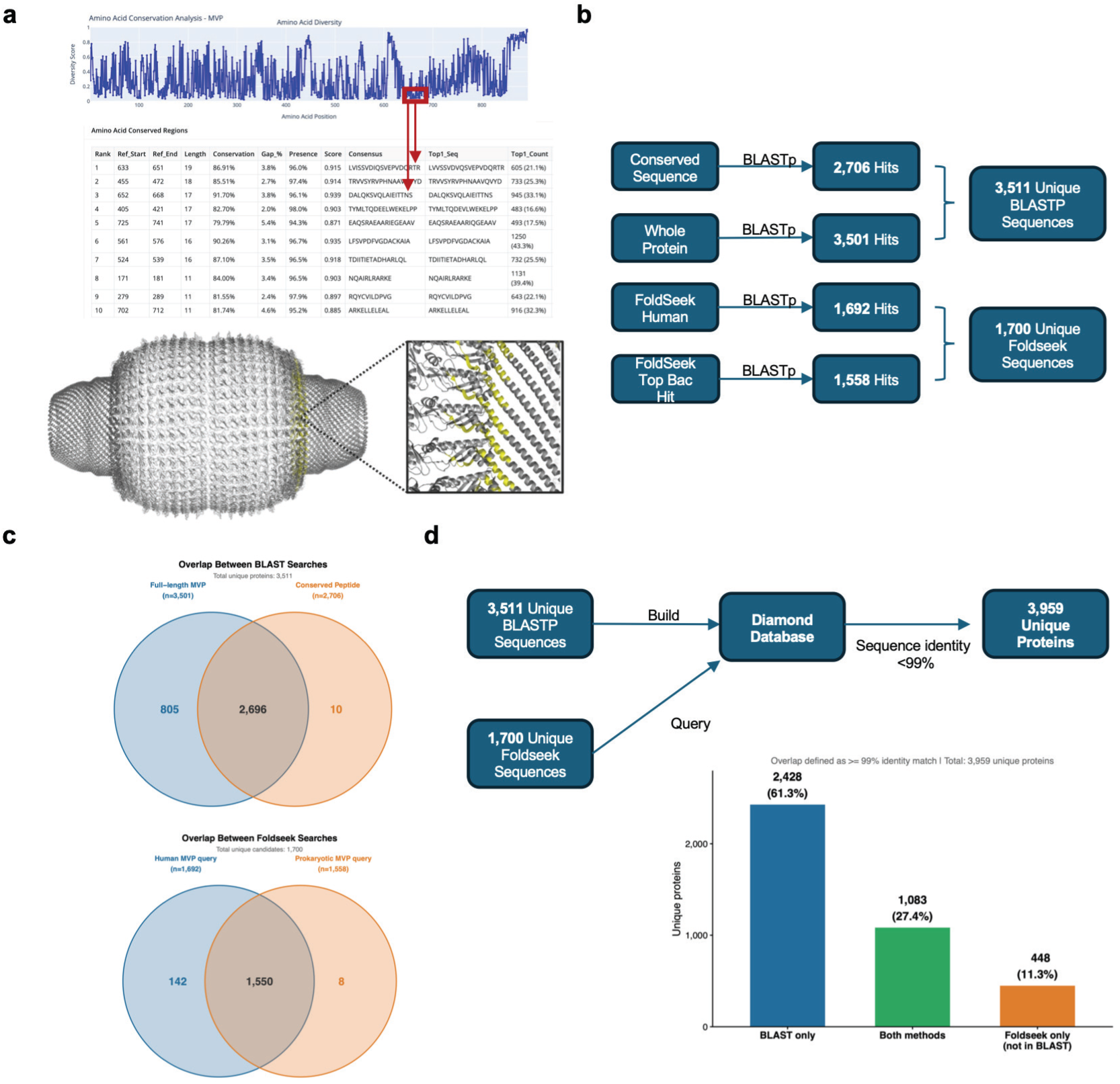
Bioinformatic analysis workflow to identify MVP homologs across the three domains of life. **a** From NCBI (Date Accessed: Jun 2, 2026), 3352 nucleotide sequences were mapped to human MVP, and a consensus sequence was identified (red box). Consensus sequence is highlighted on the human vault structure (PDB: 9BW5) in yellow. **b** Using the conserved-peptide and whole-protein sequences as BLASTp queries, we obtained 2,706 and 3,501 homologous sequences across the three domains. We further applied Foldseek structural searches with the human and prokaryotic MVP structures, recovering 1,692 and 1,558 candidates. After merging both approaches and filtering partial sequences, 3,959 unique protein sequences remained. **c** Overlap between the two BLASTp searches: the full-length MVP query (n=3,501) and the conserved-peptide query (n=2,706) shared 2,696 hits and together recovered 3,511 unique sequences (805 unique to full-length, 10 unique to the conserved peptide); the two Foldseek structural searches, the human MVP query (n=1,692) and the prokaryotic MVP query (n=1,558), shared 1,550 hits and together recovered 1,700 unique candidates (142 unique to human, 8 unique to prokaryotes). **d** The sequence- and structure-based sets were merged non-redundantly by building a DIAMOND database from the 3,511 unique BLAST sequences and querying it with the 1,700 unique Foldseek sequences, collapsing matches at >=99% identity to yield 3,959 unique MVP proteins: 2,428 (61.3%) recovered by BLAST only, 1,083 (27.4%) by both methods, and 448 (11.3%) by Foldseek only.

**Extended Data Fig. 2:**
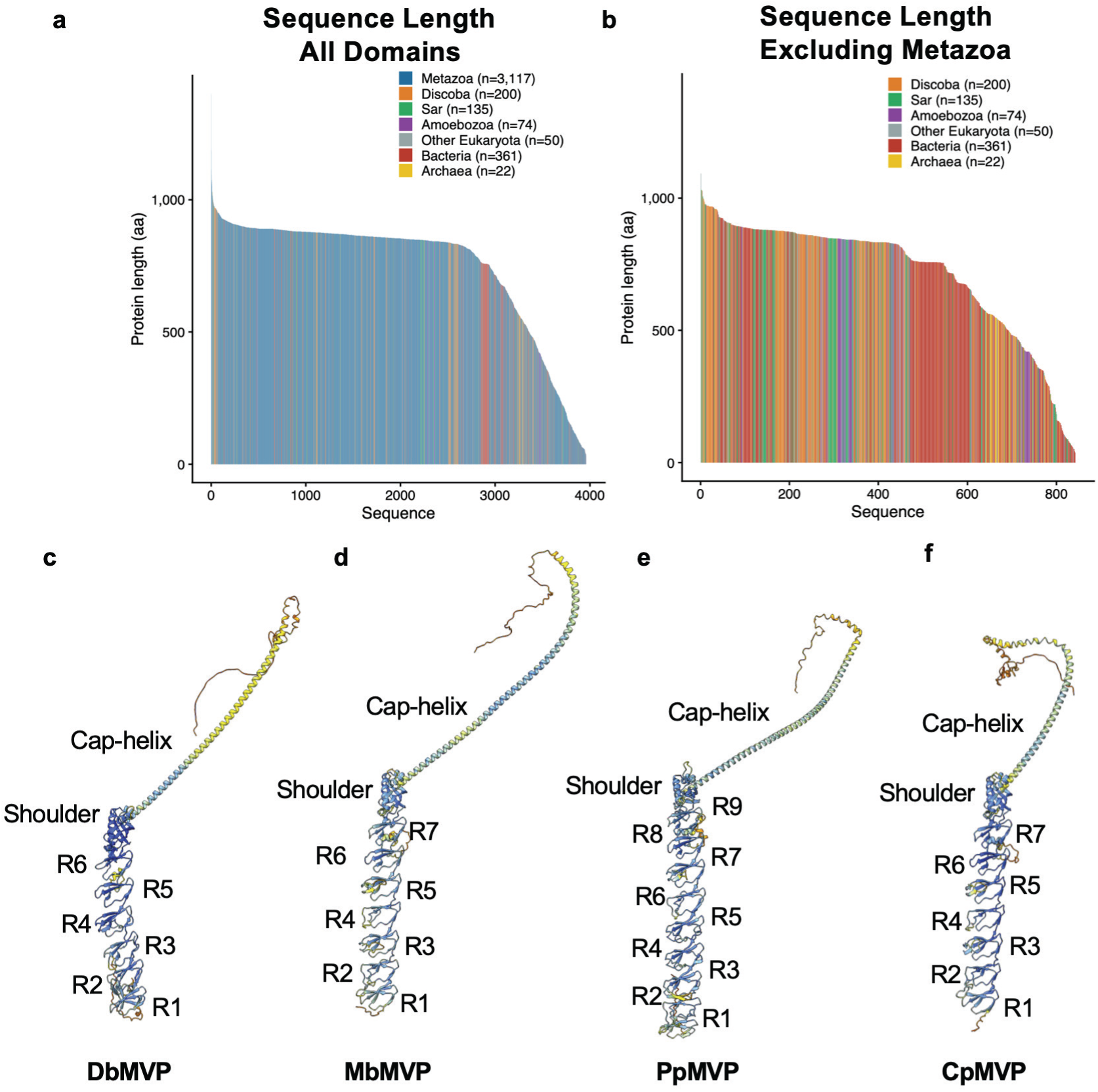
Length distribution and AlphaFold predictive modeling reveal that bacterial MVP homologs likely have similar features to mammalian MVP. **a-b** Sequence length of all BlastP hits colored by supergroups. **a** Eukaryotic MVPs are tightly conserved at near-human length. With fusions removed, the homologs span 31–1,400 aa with a median of 854 aa, closely matching human (893 aa). Metazoan homologs (n = 3,117) cluster around the human value (median 858 aa; 73% between 800 and 950 aa). The other major eukaryotic lineages are equally conserved — Discoba (median 841 aa, n = 200), Sar (847 aa, n = 135), Amoebozoa (829 aa, n = 74). **b** Prokaryotic MVPs are shorter and more variable. Bacterial homologs (n = 361) have a median length of 758 aa, and 249 of 383 prokaryotes (65%) are shorter than 800 aa. Archaeal MVPs are the most compact (n = 22; median 543 aa; most between 470 and 570 aa) **c-f** AlphaFold predictive models for **c** DbMVP, **d** MbMVP, **e** PpMVP, and **f** CpMVP. DbMVP, MbMVP, and PpMVP were from the AlphaFold protein structure database (https://alphafold.ebi.ac.uk/) whereas CpMVP was predicted using AlphaFold2. Predicted structures are colored by pLDDT score where model confidence is: Very High (Dark blue, pLDDT>90), High (Light blue, pLDDT >70), Low (Yellow, pLDDT >50), and Very low (Orange, pLDDT <50).

**Extended Data Fig. 3:**
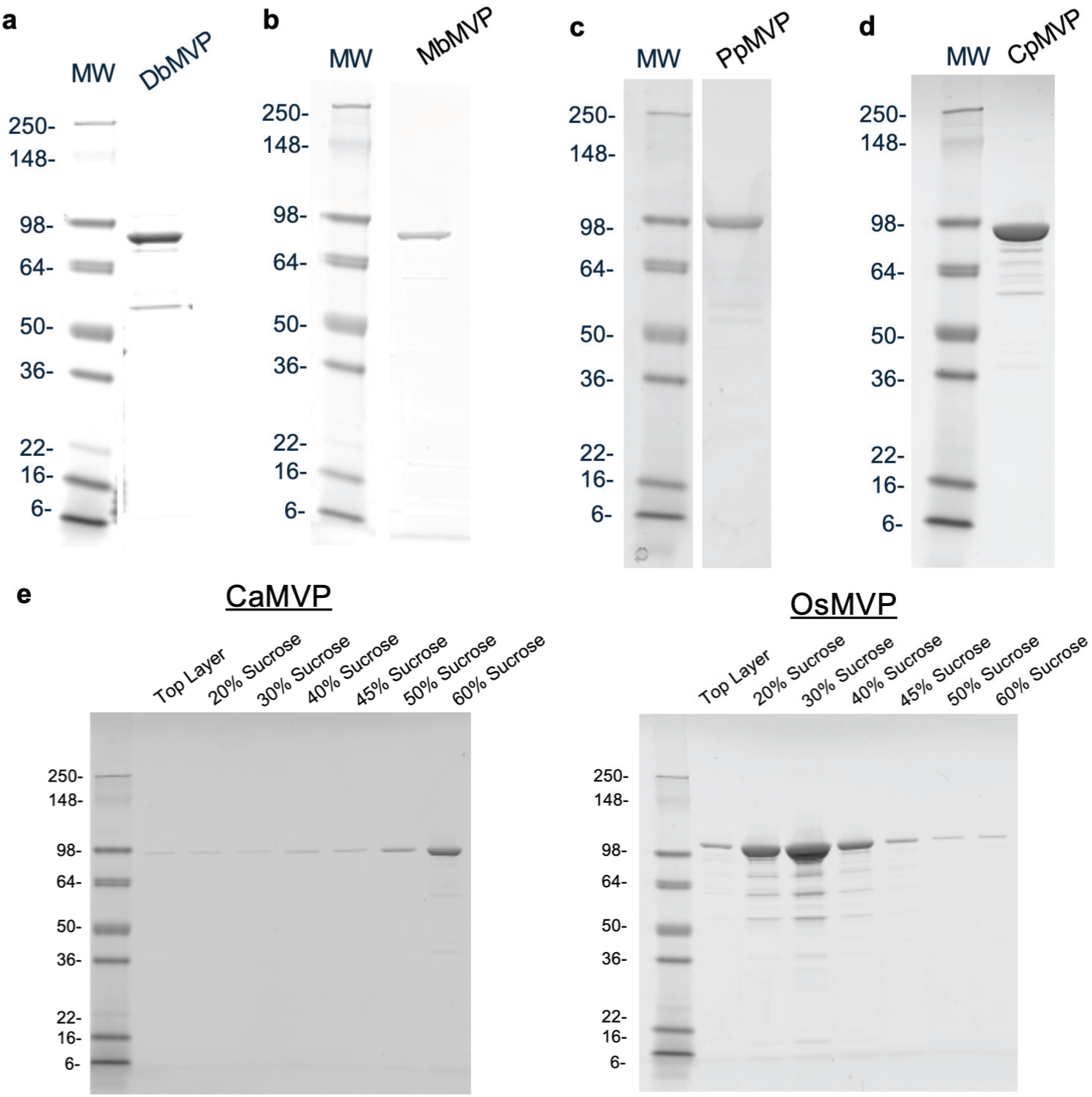
SDS-PAGE of purified proteins utilized in this study. **a-c** SDS-PAGE post-affinity purification for **a** DbMVP-6xHis, **b** MbMVP-TEV-StrepII, **c** PpMVP-TEV-StrepII, and **d** CpMVP-TEV-StrepII. A total of 2 μg of protein was loaded in each lane. e SDS-PAGE of equal volumes from each fraction on a discontinuous sucrose gradient post-affinity chromatography for CaMVP-TEV-StrepII and OsMVP-TEV-StrepII.

**Extended Data Fig. 4:**
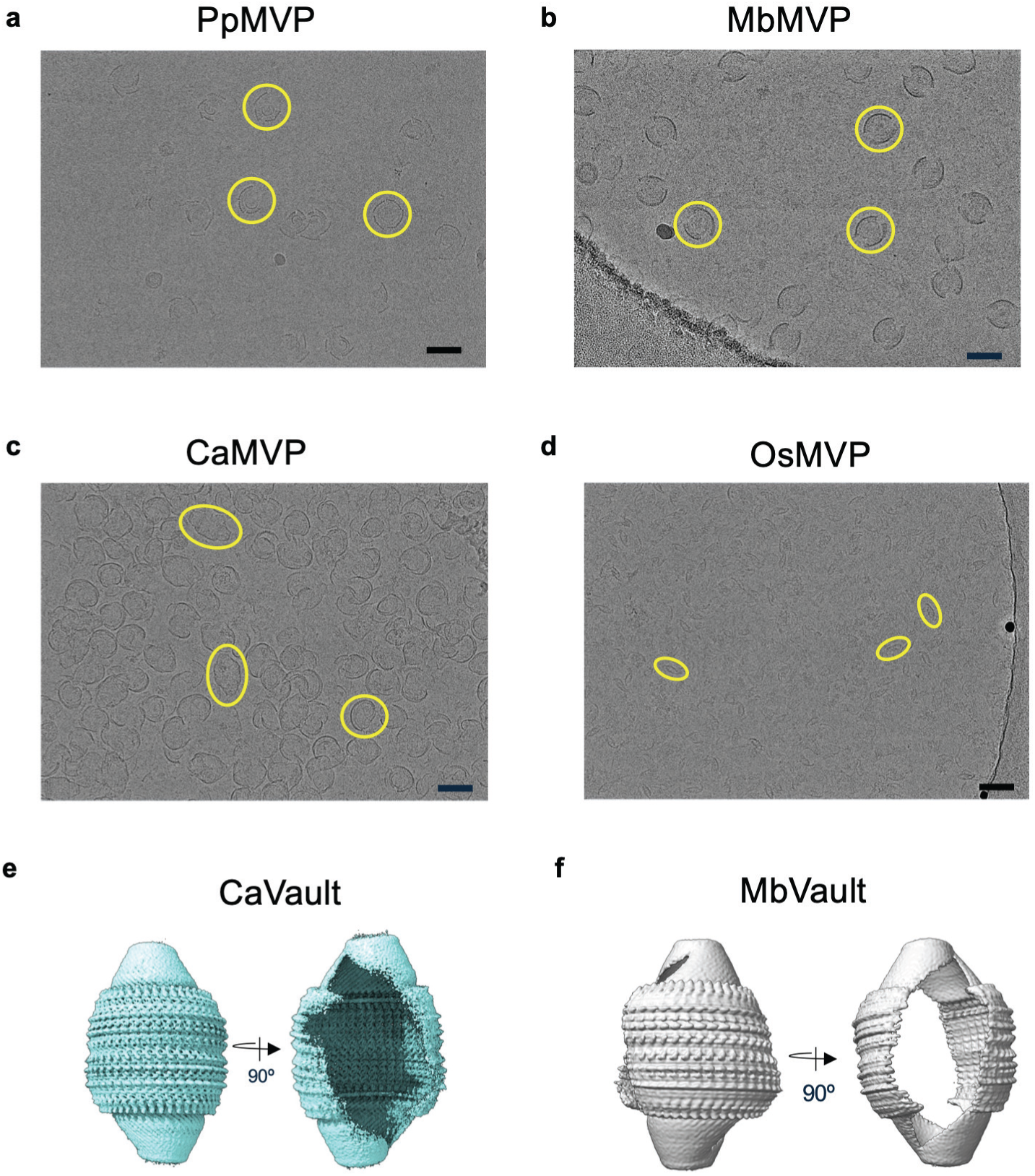
Select bacterial MVP homologs form fractured vault-like particles. **a-d** Micrographs of purified MVP homologs PpMVP (a), MbMVP (b), CaMVP (c), and OsMVP (d). While all images were acquired from proteins purified from affinity chromatography, CaMVP and OsMVP were further resolved by a discontinuous sucrose gradient, and the 60% and 45% sucrose fractions were imaged, respectively. Scale bars represent 50 nm. Select particles are highlighted in yellow for emphasis. **e-f** CryoEM 3D reconstruction of MVP homologs from CaMVP (e) and MbMVP (f). Reconstructions depict fractured vault-like structures.

**Extended Data Fig. 5:**
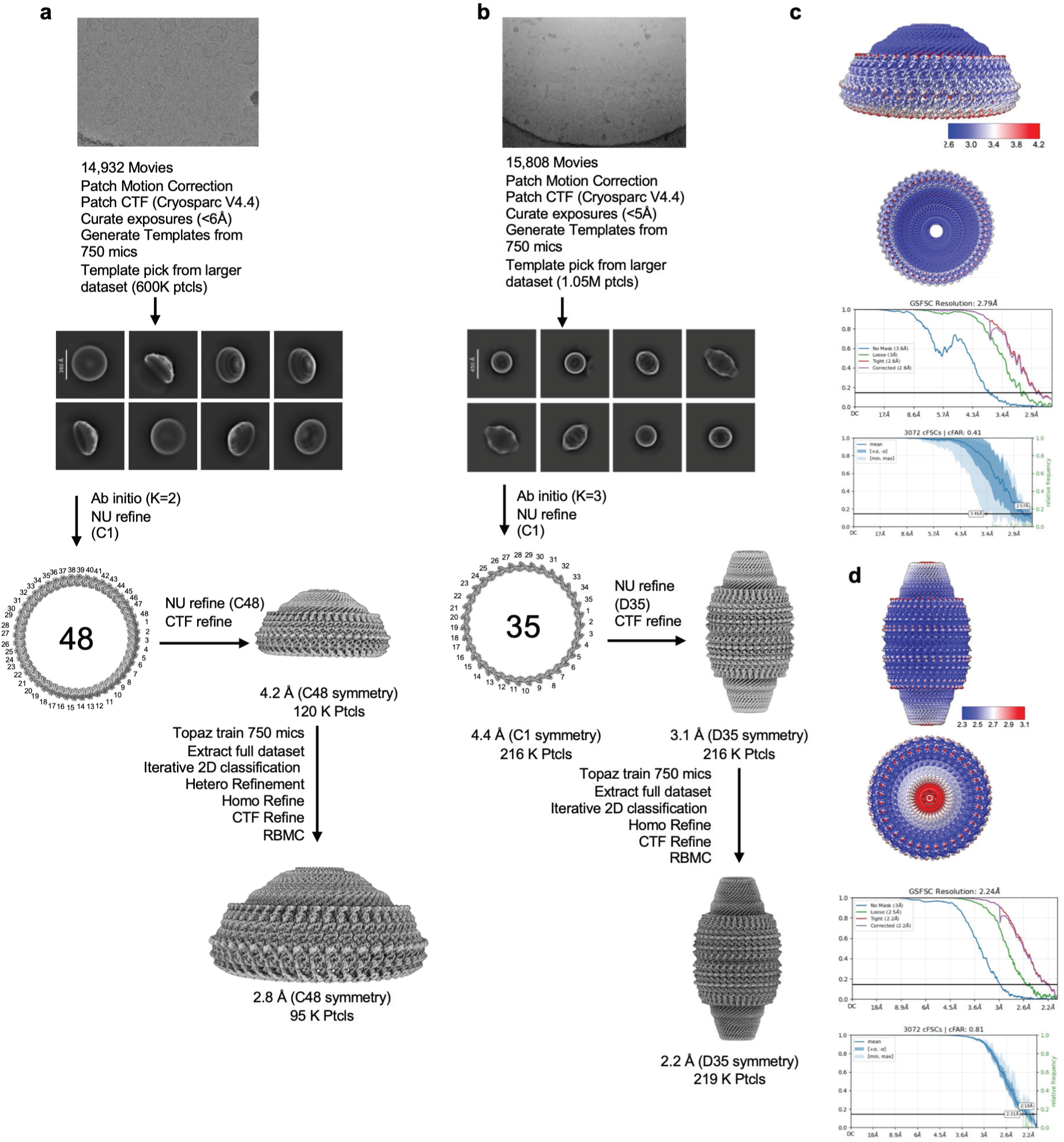
CryoEM processing of DbVault and CpVault to obtain overall structures. **a,b** CryoEM data processing pipeline for DbVault (**a**) and CpVault (**b**). Only representative 2D class averages are shown. **c,d** CryoEM map filtered and colored according to local resolution in Å, gold standard FSC curve, and conical FSC curve as produced by CryoSPARC for DbVault (**c**) and CpVault (**d**).

**Extended Data Fig. 6:**
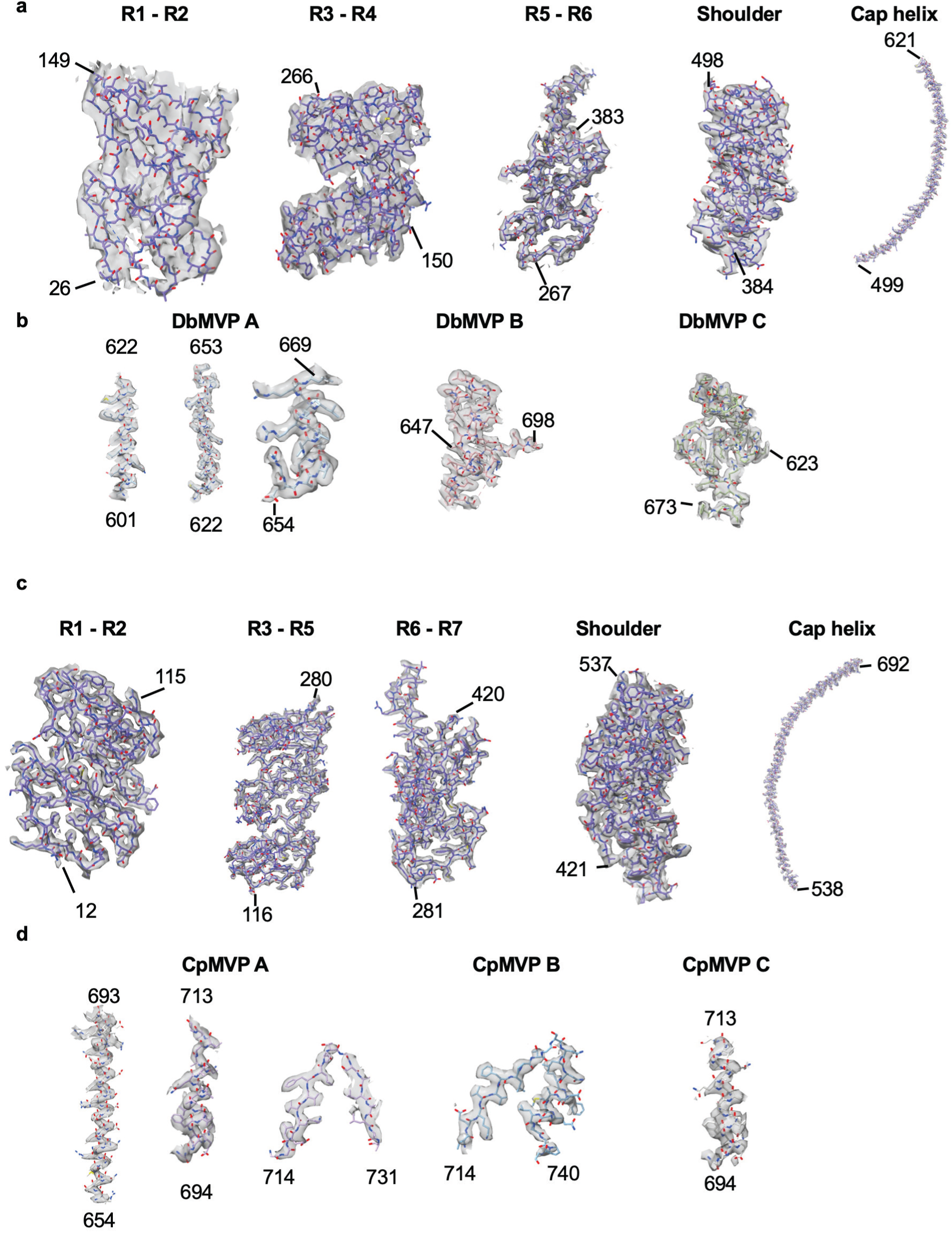
Map model fit of DbVault and CpVault. **a-d** Map to model fits of DbVault barrel (**a**), DbVault cap (**b**), CpVault barrel (**c**) and CpVault cap (**d**). C- and N-termini are labeled with residue numbers.

**Extended Data Fig. 7:**
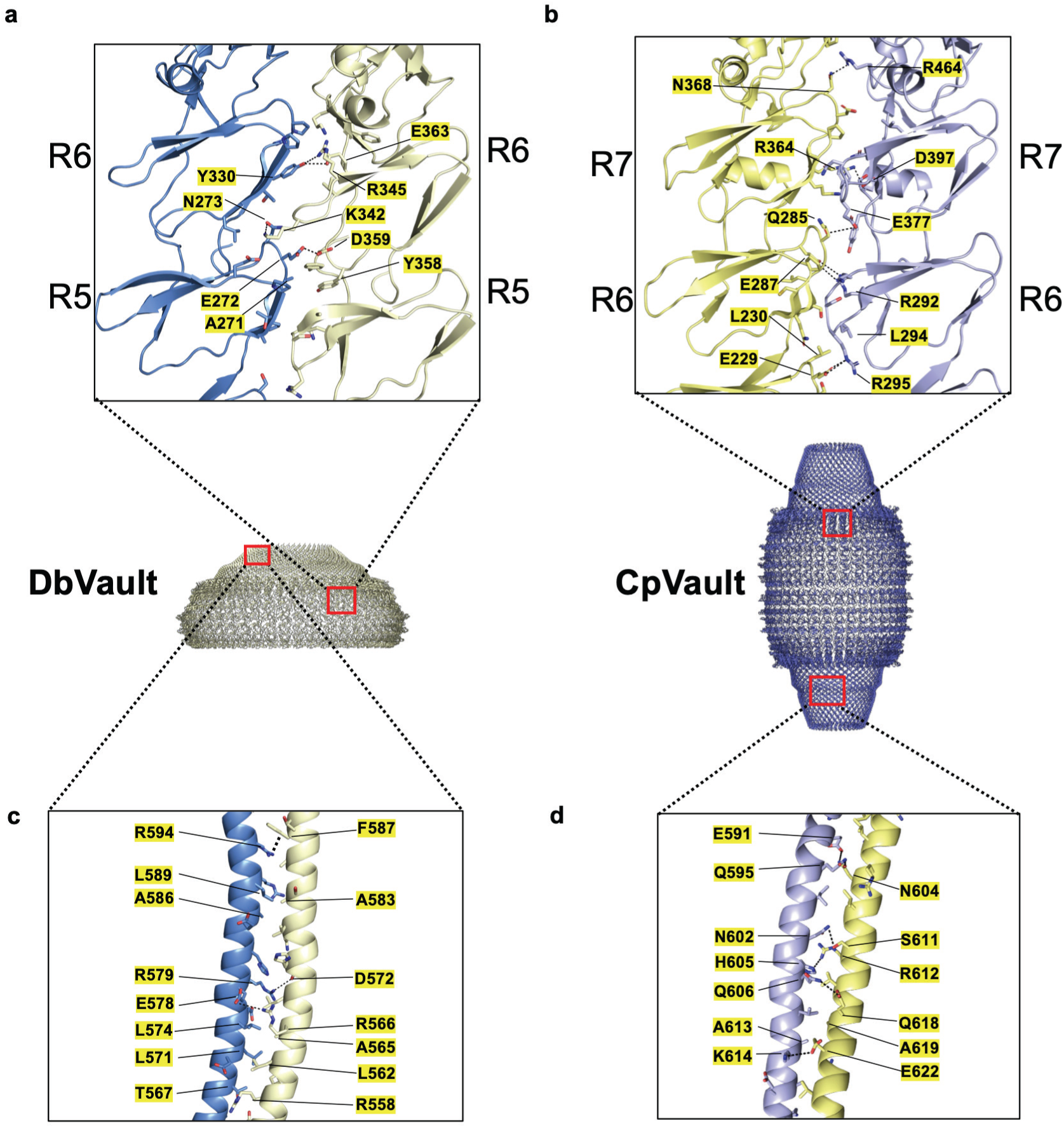
Interchain interactions within the barrel and cap-helix of DbVault and CpVault. **a-d** All residues with atoms within 4.5Å or less of the adjacent chain with selected interactions highlighted within R6 and R5 of DbVault (**a**), R7 and R6 of CpVault (**b**), the cap-helix of DbVault (**c**), and the cap-helix of CpVault (**d**).

**Extended Data Fig. 8:**
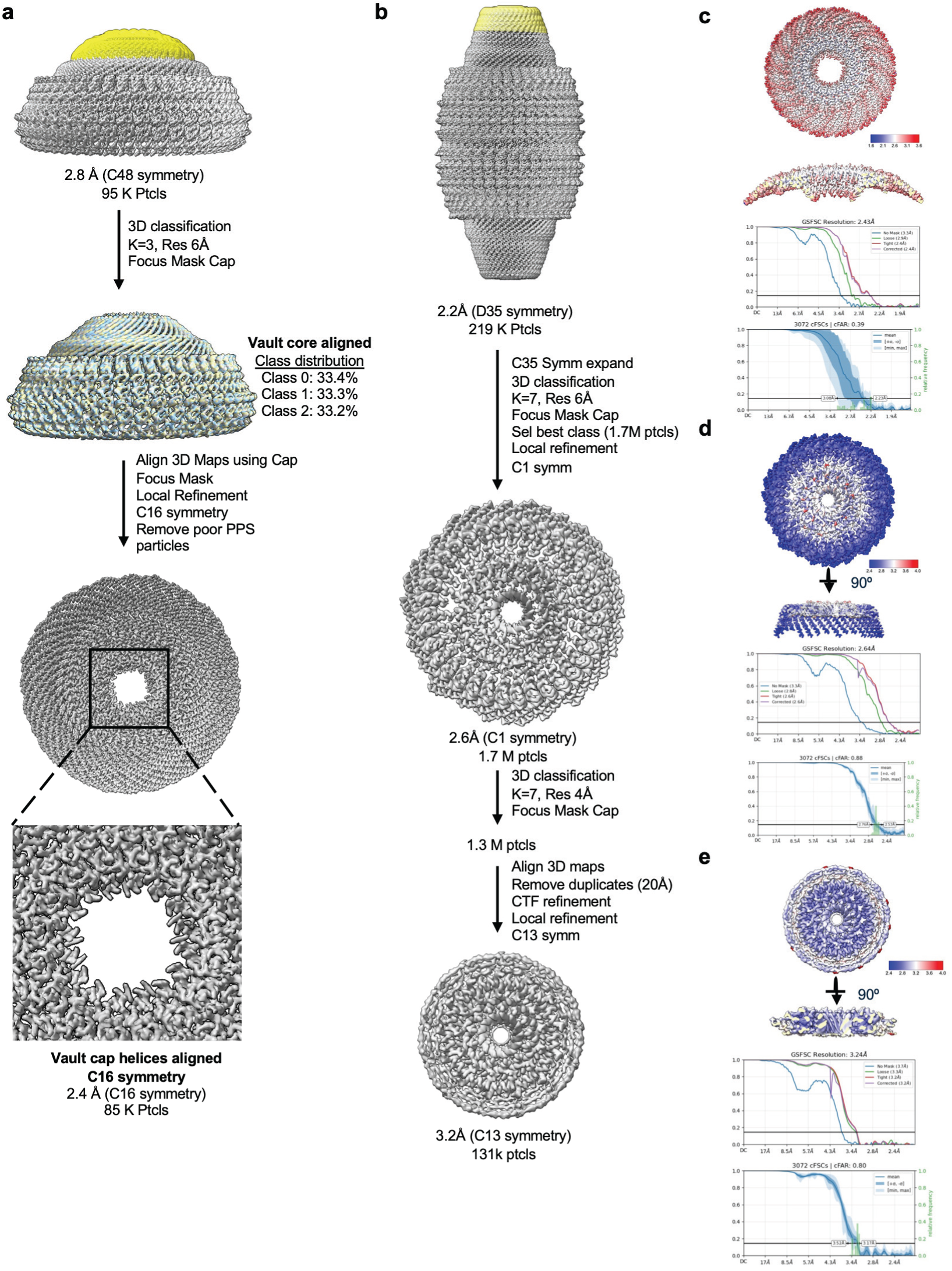
Cap-focused cryoEM processing of DbVault and CpVault. **a,b** CryoEM data processing pipeline for the DbVault cap (**a**) and CpVault cap (**b**). **c-e** CryoEM map filtered and colored according to local resolution, gold standard FSC curve, and conical FSC curve as produced by CryoSPARC for (**c**) DbVault, (**d**) CpVault (C1), and (**e**) CpVault (C13). See methods and materials for further detail.

**Extended Data Fig. 9:**
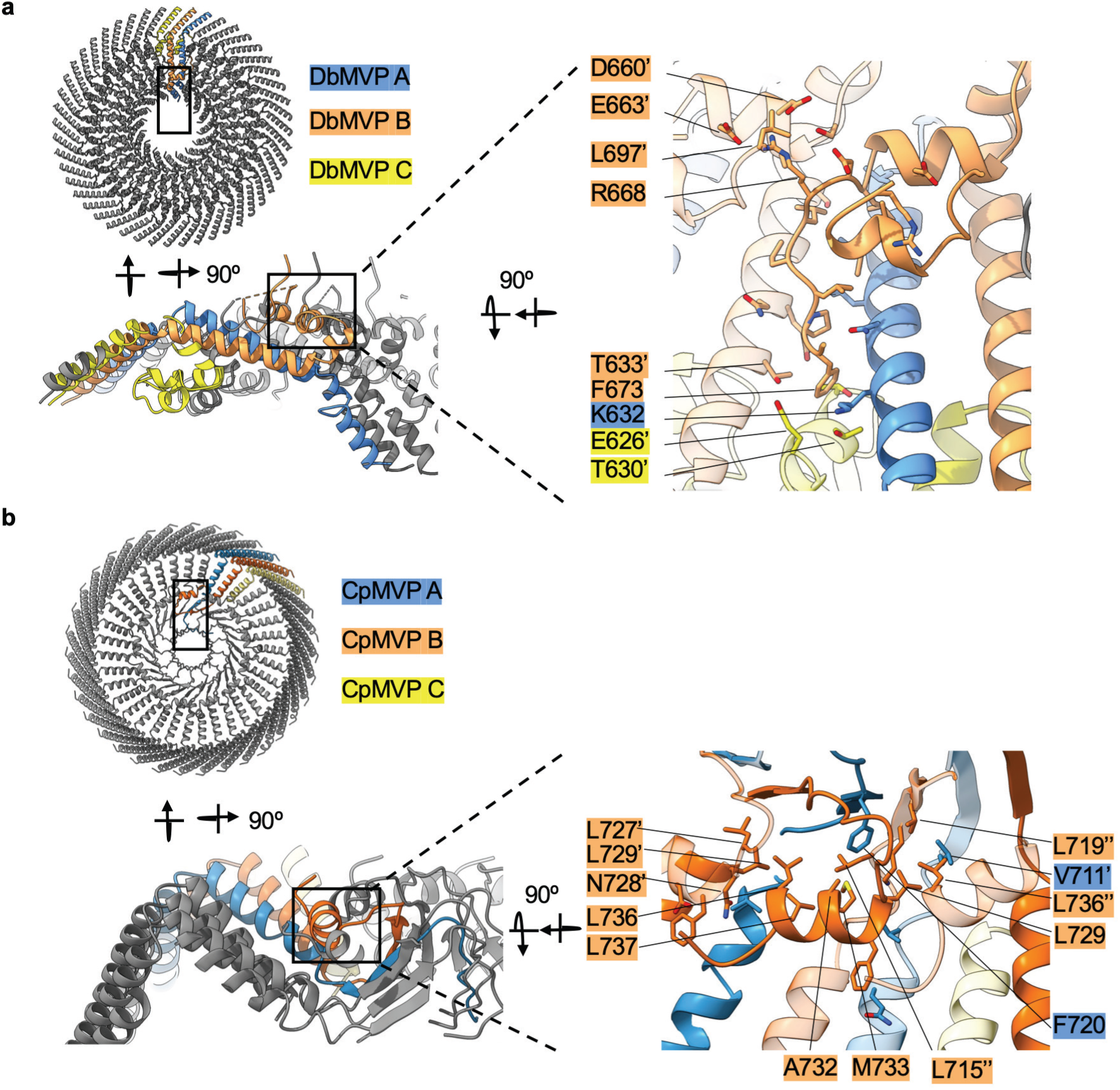
MVP B interprotomer interaction analyses in DbVault and CpVault caps. **a** Interaction network of DbMVP B and adjacent MVP protomers. **b** Interaction network of CpMVP B and adjacent MVP protomers. Monomers are color coded as they appear throughout the manuscript. Residues from neighboring asymmetric units are labeled with apostrophes.

**Extended Data Fig. 10:**
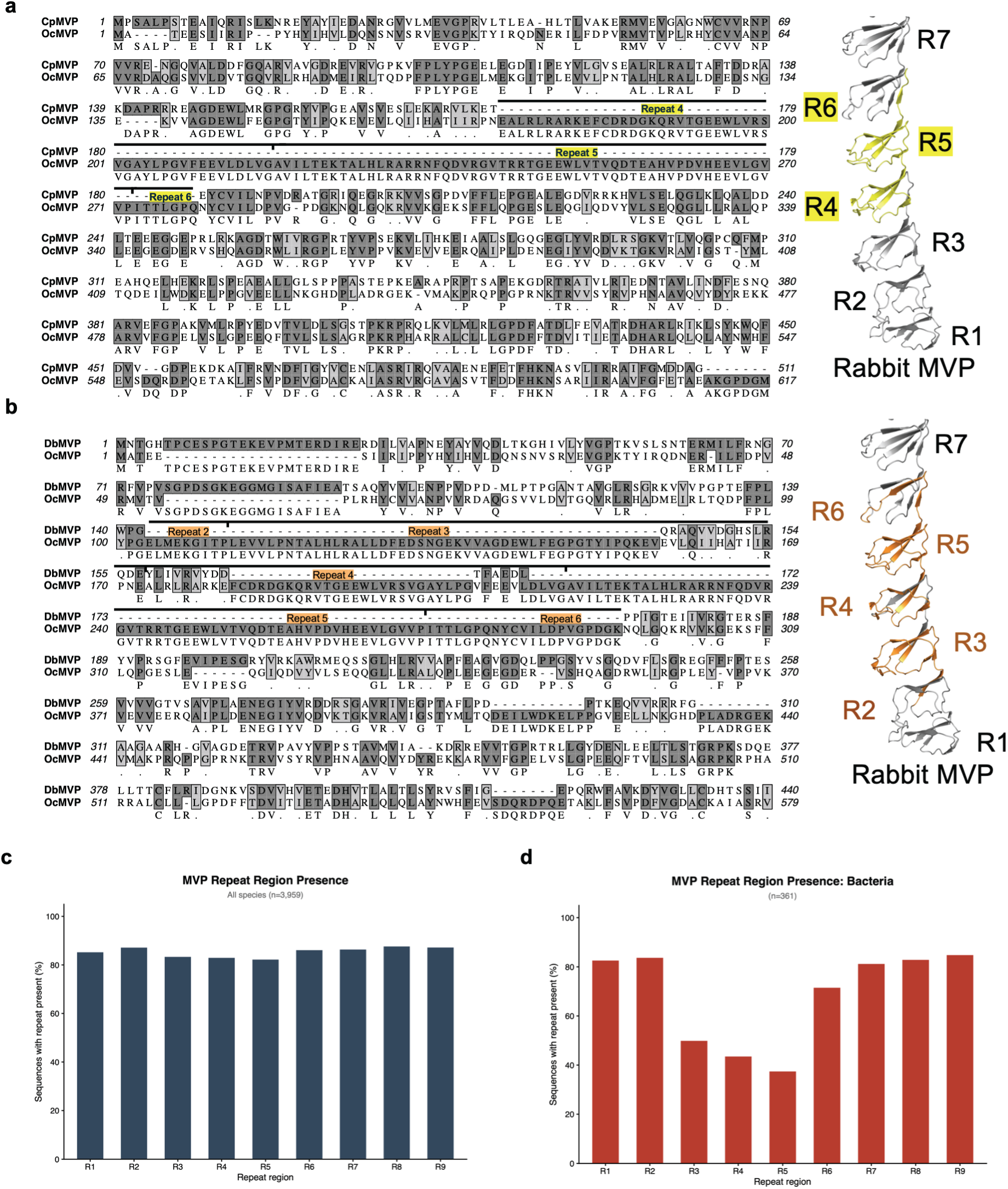
Sequence analysis indicates that R3-R5 exhibit low conservation in bacterial homologs. **a,b** Sequence alignment between of CpMVP (**a**) and DbMVP (**b**) with rabbit MVP (OcMVP), with significant gaps highlighted within the structure of rabbit MVP (PDB: 9Z4W). Sequences were aligned using T-coffee alignment (Alignment performed using MacVector: Myers-Miller, Gap open penalty -50, Gap extension penalty -5). Downward tic marks represent the bounds of each repeat domain of rabbit MVP with major gaps in the alignment. **c, d** Average percent identity of repeat domains from all species (**c**) and bacteria (**d**) identified from BlastP hits.

## References

1 Kedersha, N. L. & Rome, L. H. Isolation and characterization of a novel ribonucleoprotein particle: large structures contain a single species of small RNA. J Cell Biol 103, 699–709 (1986). 10.1083/jcb.103.3.699

2 Kedersha, N. L., Heuser, J. E., Chugani, D. C. & Rome, L. H. Vaults. III. Vault ribonucleoprotein particles open into flower-like structures with octagonal symmetry. Journal of Cell Biology 112, 225–235 (1991). 10.1083/jcb.112.2.225

3 Lodwick, J. E. et al. Structural insights into the roles of PARP4 and NAD+ binding in the human vault cage. Nature Communications 16 (2025). 10.1038/s41467-025-61981-x

4 Lapenta, F. et al. Structural flexibility of the human vault particle revealed by high-resolution cryo-EM and molecular dynamics simulations. Nature Communications (2026). 10.1038/s41467-026-72674-4

5 Osinski, A. et al. TIR-like NADases act in bacterial immunity and the RNA vault (openRxiv, 2026).

6 Lovestam, S. & Scheres, S. H. W. Cryo-EM structure of the vault from human brain reveals symmetry mismatch at its caps. Structure (2025). 10.1016/j.str.2025.07.014

7 Casañas, A. et al. New features of vault architecture and dynamics revealed by novel refinement using the deformable elastic network approach. Acta Crystallographica Section D Biological Crystallography 69, 1054–1061 (2013). 10.1107/s0907444913004472

8 Anderson, D. H., Kickhoefer, V. A., Sievers, S. A., Rome, L. H. & Eisenberg, D. Draft Crystal Structure of the Vault Shell at 9-Å Resolution. PLoS Biology 5, e318 (2007). 10.1371/journal.pbio.0050318

9 Ding, K. et al. Solution Structures of Engineered Vault Particles. Structure 26, 619–626.e613 (2018). 10.1016/j.str.2018.02.014

10 Tanaka, H. et al. The Structure of Rat Liver Vault at 3.5 Angstrom Resolution. Science 323, 384–388 (2009). 10.1126/science.1164975

11 Guerra, P. et al. Symmetry disruption commits vault particles to disassembly. Science Advances 8, eabj7795 (2022). doi:10.1126/sciadv.abj7795

12 Li, H., Vallese, F. & Clarke, O. B. The vault particle is enclosed by a *C*13-symmetric cap with a positively charged exterior. Science Advances 12 (2026). 10.1126/sciadv.adz6794

13 Gao, J. et al. Structures of human organellar SPFH protein complexes. Nature Communications 16 (2025). 10.1038/s41467-025-65078-3

14 Mrazek, J. et al. Polyribosomes Are Molecular 3D Nanoprinters That Orchestrate the Assembly of Vault Particles. ACS Nano 8, 11552–11559 (2014). 10.1021/nn504778h

15 Goldsmith, L. E., Yu, M., Rome, L. H. & Monbouquette, H. G. Vault nanocapsule dissociation into halves triggered at low pH. Biochemistry 46, 2865–2875 (2007). 10.1021/bi0606243

16 Yang, J. et al. Vaults are dynamically unconstrained cytoplasmic nanoparticles capable of half vault exchange. ACS Nano 4, 7229–7240 (2010). 10.1021/nn102051r

17 Carter, S. D., Mamede, J. I., Hope, T. J. & Jensen, G. J. Correlated cryogenic fluorescence microscopy and electron cryo-tomography shows that exogenous TRIM5α can form hexagonal lattices or autophagy aggregates in vivo. Proceedings of the National Academy of Sciences 117, 29702–29711 (2020). 10.1073/pnas.1920323117

18 Yu, Z. et al. PTEN Associates with the Vault Particles in HeLa Cells. Journal of Biological Chemistry 277, 40247–40252 (2002). 10.1074/jbc.m207608200

19 Yi, C. et al. Major Vault Protein, in Concert with Constitutively Photomorphogenic 1, Negatively Regulates c-Jun–Mediated Activator Protein 1 Transcription in Mammalian Cells. Cancer Research 65, 5835–5840 (2005). 10.1158/0008-5472.can-05-0423

20 Kim, E. et al. Crosstalk between Src and major vault protein in epidermal growth factor- dependent cell signalling. The FEBS Journal 273, 793–804 (2006). 10.1111/j.1742-4658.2006.05112.x

21 Steiner, E. et al. The major vault protein is responsive to and interferes with interferon-γ-mediated STAT1 signals. Journal of Cell Science 119, 459–469 (2006). 10.1242/jcs.02773

22 Wang, W., Xiong, L., Wang, P., Wang, F. & Ma, Q. Major vault protein plays important roles in viral infection. IUBMB Life 72, 624–631 (2020). 10.1002/iub.2200

23 Liu, S. et al. Human hepatitis B virus surface and e antigens inhibit major vault protein signaling in interferon induction pathways. Journal of Hepatology 62, 1015–1023 (2015). 10.1016/j.jhep.2014.11.035

24 Peng, N. et al. Inducible Major Vault Protein Plays a Pivotal Role in Double-Stranded RNA–or Virus-Induced Proinflammatory Response. The Journal of Immunology 196, 2753–2766 (2016). 10.4049/jimmunol.1501481

25 Liu, S. et al. Major vault protein: a virus-induced host factor against viral replication through the induction of type-I interferon. Hepatology 56, 57–66 (2012). 10.1002/hep.25642

26 Eichenmüller, B. et al. Vaults bind directly to microtubules via their caps and not their barrels. Cell Motility 56, 225–236 (2003). 10.1002/cm.10147

27 van Zon, A. et al. Vault mobility depends in part on microtubules and vaults can be recruited to the nuclear envelope. Exp Cell Res 312, 245–255 (2006). 10.1016/j.yexcr.2005.10.016

28 Maniatis, A. et al. Vault Particles in Cancer Progression, Multidrug Resistance, and Drug Delivery: Current Insights and Future Applications. International Journal of Molecular Sciences 26, 1562 (2025). 10.3390/ijms26041562

29 Rome, L. H. & Kickhoefer, V. A. Development of the Vault Particle as a Platform Technology. ACS Nano 7, 889–902 (2013). 10.1021/nn3052082

30 Wang, M., Abad, D., Kickhoefer, V. A., Rome, L. H. & Mahendra, S. Vault Nanoparticles Packaged with Enzymes as an Efficient Pollutant Biodegradation Technology. ACS Nano 9, 10931–10940 (2015). 10.1021/acsnano.5b04073

31 Voth, B. L. et al. Intratumor injection of CCL21-coupled vault nanoparticles is associated with reduction in tumor volume in an in vivo model of glioma. Journal of Neuro-Oncology 147, 599–605 (2020). 10.1007/s11060-020-03479-8

32 Buehler, D. C. et al. Bioengineered Vaults: Self-Assembling Protein Shell–Lipophilic Core Nanoparticles for Drug Delivery. ACS Nano 8, 7723–7732 (2014). 10.1021/nn5002694

33 Kar, U. K. et al. Vault Nanocapsules as Adjuvants Favor Cell-Mediated over Antibody-Mediated Immune Responses following Immunization of Mice. PLoS ONE 7, e38553 (2012). 10.1371/journal.pone.0038553

34 Kar, U. K. et al. Novel CCL21-Vault Nanocapsule Intratumoral Delivery Inhibits Lung Cancer Growth. PLoS ONE 6, e18758 (2011). 10.1371/journal.pone.0018758

35 Daly, T. K., Sutherland-Smith, A. J. & Penny, D. In Silico Resurrection of the Major Vault Protein Suggests It Is Ancestral in Modern Eukaryotes. Genome Biology and Evolution 5, 1567–1583 (2013). 10.1093/gbe/evt113

36 van Zon, A., Mossink, M. H., Scheper, R. J., Sonneveld, P. & Wiemer, E. A. The vault complex. Cell Mol Life Sci 60, 1828–1837 (2003). 10.1007/s00018-003-3030-y

37 González Álamos, M. Deciphering the structure and dynamics of the vault particles, Universitat de Barcelona, (2023).

38 Geißler, K. et al. The vault associates with membranes in situ. Nature Communications 17 (2026). 10.1038/s41467-026-71837-7

39 Vasu, S. K. & Rome, L. H. Dictyostelium Vaults: Disruption of the Major Proteins Reveals Growth and Morphological Defects and Uncovers a New Associated Protein. Journal of Biological Chemistry 270, 16588–16594 (1995). 10.1074/jbc.270.28.16588

40 Zavrelova, A. et al. The trypanosome vault particle is composed of multiple major vault protein paralogs and harbors vault RNA. J Biol Chem 301, 110706 (2025). 10.1016/j.jbc.2025.110706

41 Daly, T. K., Sutherland-Smith, A. J. & Penny, D. Beyond BLASTing: Tertiary and Quaternary Structure Analysis Helps Identify Major Vault Proteins. Genome Biology and Evolution 5, 217–232 (2013). 10.1093/gbe/evs135

42 Shaik, A. A. The unique phylogenetic distribution of vault particles reveals its functional roles, National University of Singapore, (2013).

43 Sokolskyi, T. H. Bacterial Major Vault Protein homologs shed new light on origins of the enigmatic organelle (Cold Spring Harbor Laboratory, 2019).

44 Jumper, J. et al. Highly accurate protein structure prediction with AlphaFold. Nature 596, 583–589 (2021). 10.1038/s41586-021-03819-2

45 Varadi, M. et al. AlphaFold Protein Structure Database: massively expanding the structural coverage of protein-sequence space with high-accuracy models. Nucleic Acids Research 50, D439–D444 (2022). 10.1093/nar/gkab1061

46 Varadi, M. et al. AlphaFold Protein Structure Database in 2024: providing structure coverage for over 214 million protein sequences. Nucleic Acids Research 52, D368–D375 (2024). 10.1093/nar/gkad1011

47 van Kempen, M. et al. Fast and accurate protein structure search with Foldseek. Nat Biotechnol 42, 243–246 (2024). 10.1038/s41587-023-01773-0

48 Santos, T. M., Galvão, E., Gomes, H. & Del-Bem, L.-E. Characterization and Phylogenetic Insights into the First Viral Major Vault Proteins (MVPs), Identified in Bacteriophages from the Human Gut (Cold Spring Harbor Laboratory, 2023).

49 Goddard, T. D. et al. UCSF ChimeraX: Meeting modern challenges in visualization and analysis. Protein Sci 27, 14–25 (2018). 10.1002/pro.3235

50 Meng, E. C. et al. UCSF ChimeraX: Tools for structure building and analysis. Protein Science 32, e4792 (2023). 10.1002/pro.4792

51 Pettersen, E. F. et al. UCSF ChimeraX: Structure visualization for researchers, educators, and developers. Protein Sci 30, 70–82 (2021). 10.1002/pro.3943

52 Tan, K. A. et al. Cryo-EM structure of the SPFH-NfeD family protein complex QmcA-YbbJ. Structure 32, 1603–1610.e1603 (2024). 10.1016/j.str.2024.07.022

53 Yan, L. et al. The Erlin1/2 complex is a dynamic scaffold for membrane microdomain assembly on the endoplasmic reticulum. Molecular Cell 86, 1362–1376.e1365 (2026). 10.1016/j.molcel.2026.03.001

54 Fu, Z. & MacKinnon, R. Structure of the flotillin complex in a native membrane environment. Proceedings of the National Academy of Sciences 121, e2409334121 (2024). doi:10.1073/pnas.2409334121

55 Ma, C. et al. Structural insights into the membrane microdomain organization by SPFH family proteins. Cell Research 32, 176–189 (2022). 10.1038/s41422-021-00598-3

56 Woodward, C. L., Mendonça, L. M. & Jensen, G. J. Direct visualization of vaults within intact cells by electron cryo-tomography. Cell Mol Life Sci 72, 3401–3409 (2015). 10.1007/s00018-015-1898-y

57 Perina, D. et al. Distribution of protein poly(ADP-ribosyl)ation systems across all domains of life. DNA Repair (Amst) 23, 4–16 (2014). 10.1016/j.dnarep.2014.05.003

58 Köstlbacher, S. et al. Prediction of eukaryotic cellular complexity in Asgard archaea using structural modelling. Nature Microbiology 11, 747–758 (2026). 10.1038/s41564-026-02273-y

59 Sayers, E. W. et al. Database resources of the National Center for Biotechnology Information in 2023. Nucleic Acids Res (2022). 10.1093/nar/gkac1032

60 Edgar, R. C. MUSCLE: multiple sequence alignment with high accuracy and high throughput. Nucleic Acids Res 32, 1792–1797 (2004). 10.1093/nar/gkh340

61 Altschul, S. F., Gish, W., Miller, W., Myers, E. W. & Lipman, D. J. Basic local alignment search tool. J Mol Biol 215, 403–410 (1990). 10.1016/S0022-2836(05)80360-2

62 Buchfink, B., Reuter, K. & Drost, H. G. Sensitive protein alignments at tree-of-life scale using DIAMOND. Nat Methods 18, 366–368 (2021). 10.1038/s41592-021-01101-x

63 Katoh, K. & Standley, D. M. MAFFT: iterative refinement and additional methods. Methods Mol Biol 1079, 131–146 (2014). 10.1007/978-1-62703-646-7_8

64 Capella-Gutierrez, S., Silla-Martinez, J. M. & Gabaldon, T. trimAl: a tool for automated alignment trimming in large-scale phylogenetic analyses. Bioinformatics 25, 1972–1973 (2009). 10.1093/bioinformatics/btp348

65 Nguyen, L. T., Schmidt, H. A., von Haeseler, A. & Minh, B. Q. IQ-TREE: a fast and effective stochastic algorithm for estimating maximum-likelihood phylogenies. Mol Biol Evol 32, 268–274 (2015). 10.1093/molbev/msu300

66 Price, M. N., Dehal, P. S. & Arkin, A. P. FastTree: computing large minimum evolution trees with profiles instead of a distance matrix. Mol Biol Evol 26, 1641–1650 (2009). 10.1093/molbev/msp077

67 Mai, U., Sayyari, E. & Mirarab, S. Minimum variance rooting of phylogenetic trees and implications for species tree reconstruction. PLoS One 12, e0182238 (2017). 10.1371/journal.pone.0182238

68 Punjani, A., Rubinstein, J. L., Fleet, D. J. & Brubaker, M. A. cryoSPARC: algorithms for rapid unsupervised cryo-EM structure determination. Nature Methods 14, 290–296 (2017). 10.1038/nmeth.4169

69 Bepler, T. et al. Positive-unlabeled convolutional neural networks for particle picking in cryo-electron micrographs. Nat Methods 16, 1153–1160 (2019). 10.1038/s41592-019-0575-8

70 Liebschner, D. et al. Macromolecular structure determination using X-rays, neutrons and electrons: recent developments in *Phenix*. Acta Crystallographica Section D Structural Biology 75, 861–877 (2019). 10.1107/s2059798319011471

71 Emsley, P., Lohkamp, B., Scott, W. G. & Cowtan, K. Features and development of Coot. Acta Crystallogr D Biol Crystallogr 66, 486–501 (2010). 10.1107/s0907444910007493

72 Schrodinger, LLC. The PyMOL Molecular Graphics System, Version 1.*8* (2015).

73 Shapovalov, M. V. & Dunbrack, R. L., Jr. A smoothed backbone-dependent rotamer library for proteins derived from adaptive kernel density estimates and regressions. Structure 19, 844–858 (2011). 10.1016/j.str.2011.03.019

74 Davis, I. W. et al. MolProbity: all-atom contacts and structure validation for proteins and nucleic acids. Nucleic Acids Res 35, W375–383 (2007). 10.1093/nar/gkm216

