## Supplementary Information for "Bacterial major vault protein homologs form divergent vault structures"

**Supplementary Table 1. Conditions used for MVP homolog purification**

<table>
<tr><th>Construct</th><th>Lysis Buffer</th><th>Wash Buffer</th><th>Elution Buffer</th></tr>
<tr><td>DbMVP</td><td>50 mM Tris-HCl, pH 7.5, 75 mM NaCl, 0.75mM $MgCl_2$, 1% Triton X-305, 5% glycerol, 5mM Imidazole</td><td>50 mM Tris-HCl pH 7.5, 75 mM NaCl, 0.75mM $MgCl_2$, 5% glycerol, 5mM Imidazole</td><td>50 mM Tris-HCl, pH 7.5, 75 mM NaCl, 0.75mM $MgCl_2$, 5% glycerol, 500mM Imidazole</td></tr>
<tr><td>CpMVP, CaMVP, OsMVP</td><td>50 mM Tris-HCl, pH 7.5, 75 mM NaCl, 0.75mM $MgCl_2$, 1% Triton X-305, 5% glycerol</td><td>50 mM Tris-HCl, pH 7.5, 75 mM NaCl, 0.75mM $MgCl_2$, 5% glycerol</td><td>50 mM Tris-HCl pH 7.5, 75 mM NaCl, 0.75mM $MgCl_2$, 5% glycerol, 50mM biotin</td></tr>
<tr><td>MbMVP</td><td>100 mM Tris HCl pH 7.5, 75mM NaCl, 0.75mM $MgCl_2$, 1% Triton X-305, 1mM PMSF, 1x Halt Protease Inhibitor</td><td rowspan="2">50 mM Tris HCl pH 7.5, 75mM NaCl, 0.75mM $MgCl_2$</td><td rowspan="2">50 mM Tris HCl pH 7.5, 75mM NaCl, 0.75mM $MgCl_2$, 50mM biotin</td></tr>
<tr><td>PpMVP</td><td>50 mM Tris-HCL, pH 7.5, 75 mM NaCl, 0.75mM $MgCl_2$, 1% Triton X-305, 1mM PMSF, 1x Halt protease inhibitor</td></tr>
</table>

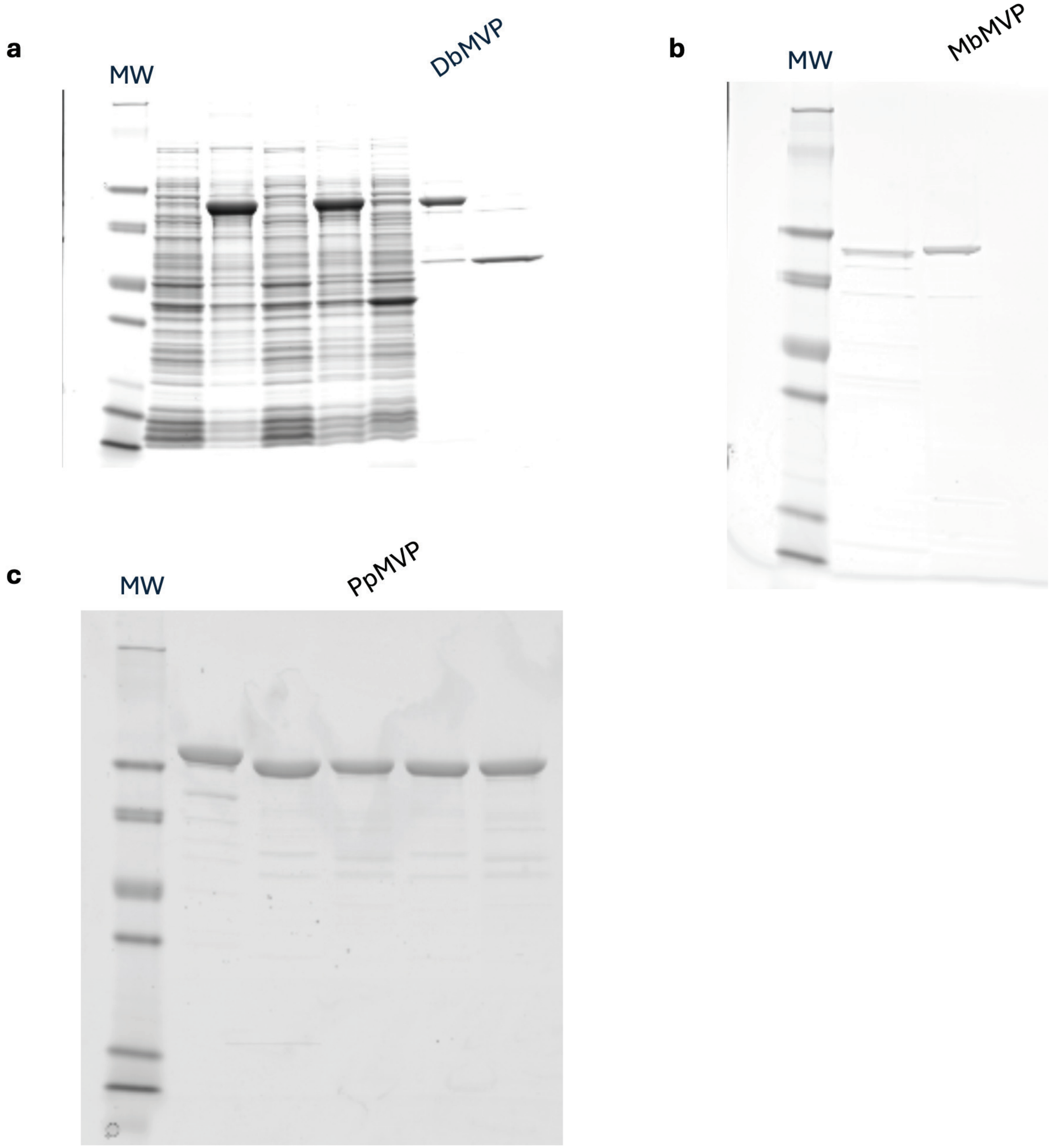


**Supplementary Fig. 1 | Uncropped Gels from Extended Data Fig. 3.**
**a-c** Uncropped SDS-PAGE post affinity purification for **a** DbMVP-6xHis, **b** MbMVP-TEV-StrepII, and **c** PpMVP-TEV-StrepII. A total of 2 µg of protein was loaded in each lane.

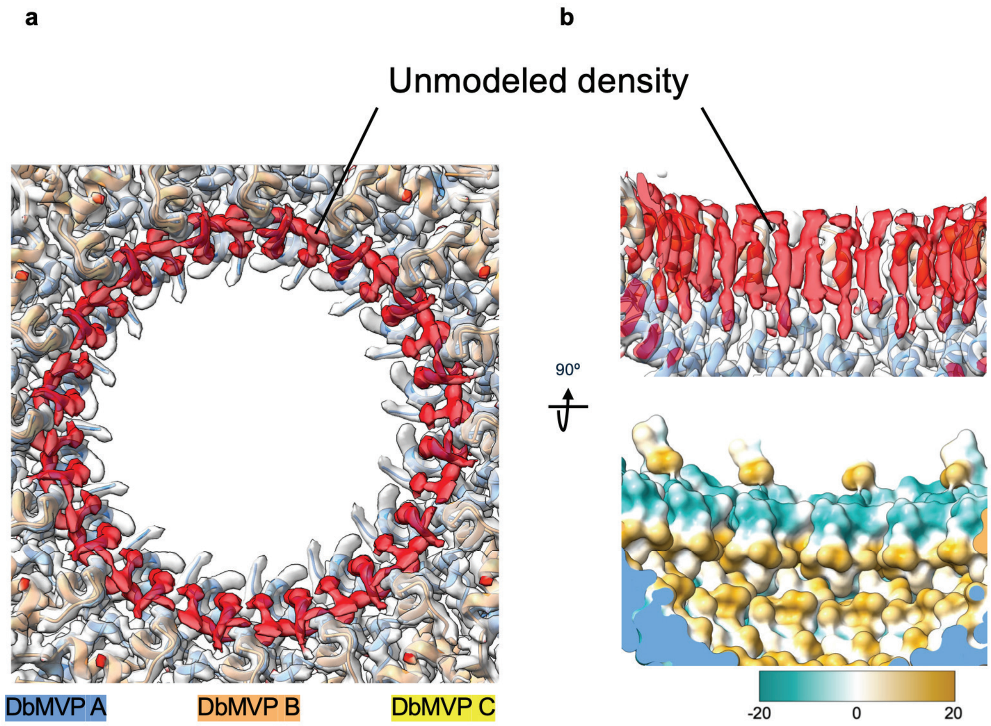


**Supplementary Fig. 2 | Unmodeled density in the upper aperture of the DbVault cap.** **a** Top-down view of the DbVault cap map colored by chain (DbMVP A,B,C) Unmodeled densities are colored red. **b** Side view of the upper aperture of the DbVault cap with map colored by chain as in panel **a** (top) and colored by hydrophobicity (arbitrary units, where teal is less hydrophobic and orange is more hydrophobic) without cryoEM map displayed (bottom).

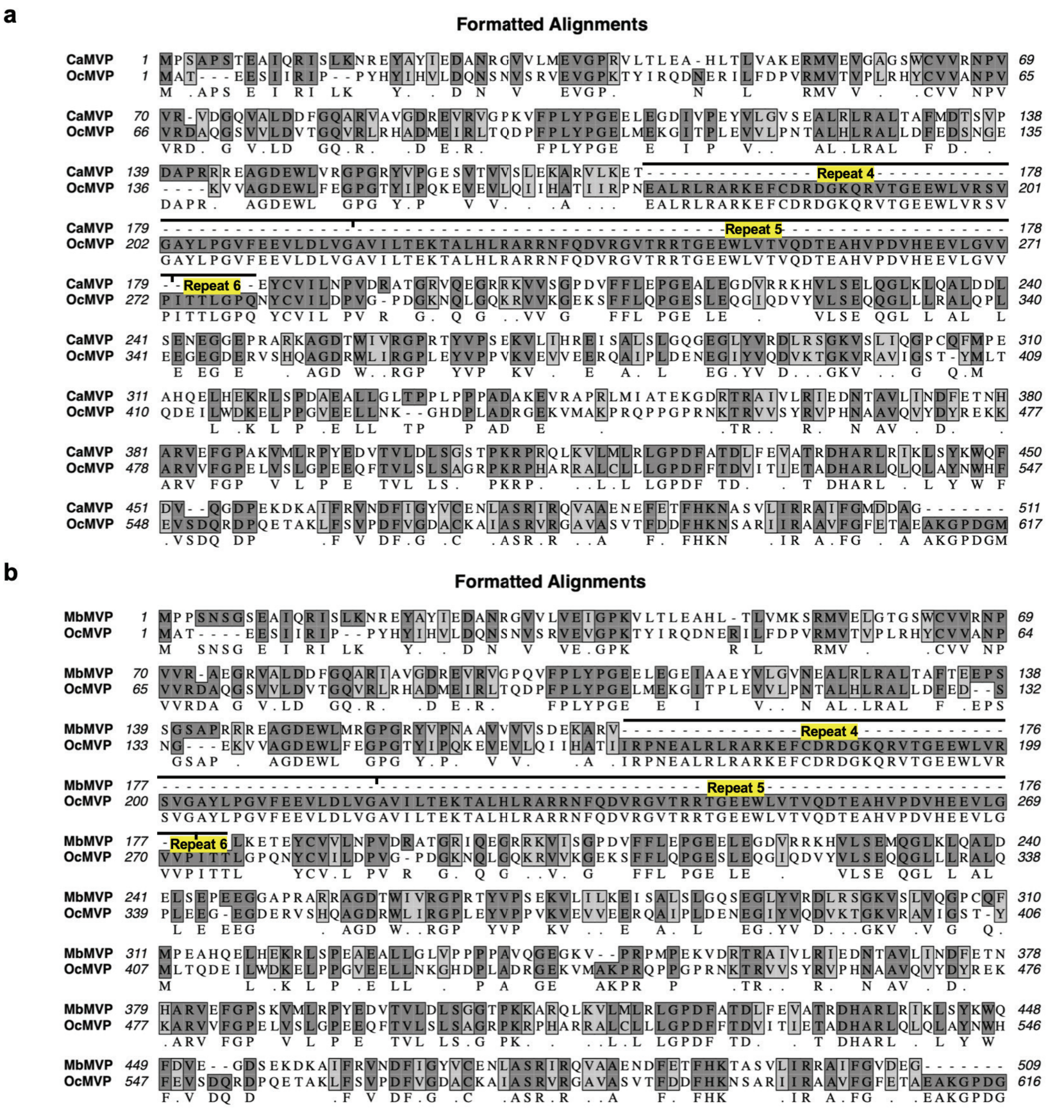


**Supplementary Fig. 3 | Sequence alignments of CaMVP and MbMVP to rabbit MVP.** **a-b** Sequence alignment of rabbit MVP (OcMVP) to CaMVP (**a**) and MbMVP (**b**). Sequences were aligned using T-coffee alignment (Alignment performed using MacVector: Myers-Miller, Gap open penalty -50, Gap extension penalty -5). Downward tic marks represent the bounds of each repeat domain of rabbit MVP with major gaps in the alignment.
